# Neuroimmune changes underscore pain-associated behaviors and disc herniations in SM/J mice

**DOI:** 10.1101/2024.02.23.581794

**Authors:** Emanuel J. Novais, Olivia K. Ottone, Eric V. Brown, Vedavathi Madhu, Victoria A. Tran, Abhijit S. Dighe, Michael D. Solga, Alexandra Manchel, Angelo C. Lepore, Makarand V. Risbud

**Affiliations:** Local Health Unit of the Litoral Alentejano, Orthopaedic Department, Santiago do Cacém, Portugal; Department of Orthopaedic Surgery, Sidney Kimmel Medical College, Thomas Jefferson University, Philadelphia, USA; Graduate Program in Cell Biology and Regenerative Medicine, Jefferson College of Life Sciences, Thomas Jefferson University, Philadelphia, USA; Faculty of Medicine, Universidade Católica Portuguesa, Portugal; Center for Interdisciplinary Research in Health, Universidade Católica Portuguesa, Portugal; Department of Neuroscience, Vickie and Jack Farber Institute for Neuroscience·, Sidney Kimmel Medical College at Thomas Jefferson University, Philadelphia, PA, USA; Department of Orthopedic Surgery, University of Virginia Health System, Charlottesville, VA, United States; Center for Public Health Genomics, University of Virginia, Charlottesville, VA, United States; Flow Cytometry Core Facility, University of Virginia, Charlottesville, VA, United States

## Abstract

There are no appropriate mouse models to study the pathophysiology of spontaneous disc herniations and associated pain pathology. We demonstrate that SM/J mice show a high incidence of age-associated lumbar disc herniations with neurovascular innervations. Transcriptomic comparisons of the SM/J annulus fibrosus with human tissues showed shared pathways related to immune cell activation and inflammation. Notably, aged SM/J mice showed increased pain sensitization and neuroinflammatory signatures associated with altered extracellular matrix regulation in the DRGs and spinal cord. There were increased T cells in the vertebral marrow, and CyTOF analysis showed increased splenic CD8^+^ T cells, nonspecific activation of CD8^+^ memory T cells, and enhanced IFN-γ production in the myeloid compartment. ScRNA-seq of PBMCs in SM/J showed more B cells, with lower proportions of T cells, monocytes, and granulocytes. This study identifies SM/J mice as a clinically-relevant model to study the pathophysiology of spontaneous disc herniations and highlights a causative axis for chronic discogenic pain with novel contributors from the primary lymphoid organs (spleen and vertebral marrow), circulation, and the nervous system.

**One-Sentence Summary:** The novel SM/J mouse model shows a neuroimmune axis drives chronic back pain, a leading cause of years lived with disability.

## Introduction

With increased human lifespan, the interest in understanding age-dependent diseases has grown(*1*). Musculoskeletal diseases, including osteoarthritis, osteoporosis, and intervertebral disc degeneration, are associated with aging, chronic pain, loss of mobility, and disability(*2*). Disc degeneration is one of the major contributors to chronic low back pain (cLBP), a top cause of years lived with disability (*3*).

While disc pathology manifests as a host of phenotypes, including fibrosis, calcification, and herniation(*4*), understanding each is hampered by the lack of appropriate animal models. Although mouse models are developed to understand these phenotypes(*5–7*), studies of herniation are limited to injury models that capture local changes after acute herniations but fail to recapitulate age-related chronic herniations(*8*). Similarly, transgenic and knockout mice have shown increased susceptibility to spontaneous herniations(*9*, *10*) and corresponding immune changes but have limited translatability to humans due to single gene contributions and other comorbidities(*11*). 85% of symptomatic human disc herniations are resorbed after 6 weeks due to neutrophil/macrophage and immune cell recruitment by the local pro-inflammatory signals(*12*). Recent studies indicate that inhibiting these early inflammatory processes delays resorption and causes a transition to a chronic pain state(*13*, *14*).

Many degenerative pathologies associated with chronic pain show increased innervation related to local expression of nerve growth factor (NGF) and protein gene product (PG) 9.5(*15*, *16*). Notably, markers such as calcitonin gene-related peptide (CGRP) and isolectin B4 (IB4) are used to discern peptidergic and non-peptidergic nociceptive neurons(*17*). While discs, facet joints, muscle, ligament, and fascia all can contribute to back pain, the sensory information is transmitted through identical circuits: local peripherical nociceptive neurons in dorsal root ganglia (DRG), the dorsal horn in the spinal cord and finally the specific sensory brain region(*18*). Furthermore, immune and glial activation in the central and peripheral nervous systems is thought to mediate the transition from acute to chronic pain(*19*)

In the present study, we provide comprehensive analyses of the pathophysiology and interdependence of spontaneous lumbar herniations and radiculopathy in SM/J mice, which are highly susceptible to age-dependent herniations(*20*). Broadly, our studies highlight the role of immune system dysregulation in chronic pain due to disc herniation and emphasize the importance of genetic background to the susceptibility of age-associated herniations.

## Results

### SM/J mice show a high incidence of age-associated lumbar disc herniations

We characterized the lumbar spinal phenotype of SM/J mice at 6-, 12-, and 20 months (M) and compared it to age-matched C57BL/6 mice. Histological analysis showed disc degeneration in SM/J at 6M, which progressed in severity at 12M and 20M, characterized by loss of NP/AF demarcation, decreased NP and AF cellularity, and increased AF clefts (Fig.1a-c’). While differences in grading distribution were smaller at 6M, there was a clear increase in higher, more degenerative grades by 20M (Fig.1d-d’). As expected, both strains demonstrated age-associated lumbar disc degeneration; however, in SM/J it was early onset and severe, evidenced by higher NP grades at 6M and NP and AF grades at 12M and 20M (Fig. 1e-e’). Strikingly, approximately 90% of SM/J mice showed herniations at multiple lumbar levels by 20M, with the highest incidence at L2-L5 (Fig. 1f-g, Suppl. Fig. 1a).

**Fig. 1:**
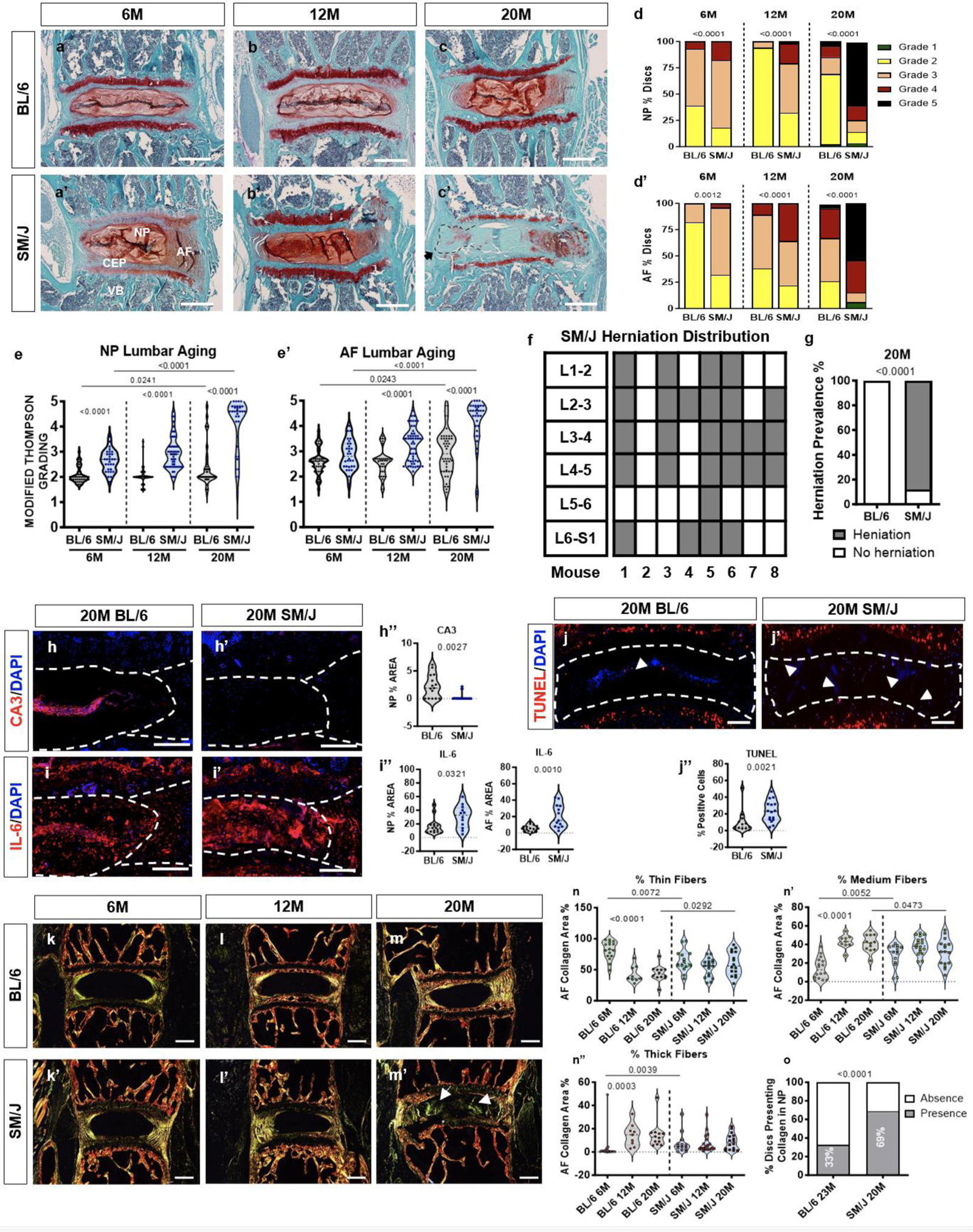
SM/J mice show high incidence of age-dependent disc herniation, accompanied by cellular deficiencies and matrix remodeling. (a-c’) Histology of the 6M, 12M, and 20M old BL/6 and SM/J lumbar discs. SM/J mice show changes in tissue architecture and cell morphology as signs of disc degeneration in the NP, AF, and CEP. (d-e’) Modified Thompson grade distributions and averages show higher average scores of degeneration in SM/J mice than BL/6 in NP and AF compartments. (f-g) Schematic showing prevalence of herniations along the lumbar spine in 20M SM/J mice. BL/6: 6M (n=5), 12M (n=7), 20M (n=11). SM/J: 6M (n=6), 12M (n=6), 20M (n=8). 4-6 lumbar discs per group were analyzed. (h-j”) Staining and abundance of key markers of (h-h”) NP cell phenotype, CA3; i-i” local inflammation, IL-6 and (j-j”) and cells viability, TUNEL. BL/6: 12M (n=5-6 mice), 20M (n=5-6). SM/J: 12M (n=5-6), 20M (n=5-6), 2-3 disc levels/animal. Scale bar = 200 µm. Scale bar = 200 µm. NP: nucleus pulposus; AF: annulus fibrosus; EP: endplate; GP: growth plate, VB: vertebral bone. (k-m’) Picrosirius Red staining and polarized microscopy of lumbar disc sections of BL/6 and SM/J discs. (n-n”) Analysis of percentage of thin (green), intermediate (yellow), and thick fibers (red). BL/6: 12M (n=5-6), 20M (n=5-6). SM/J: 12M (n=5-6), 20M (n=5-6), 2-3 disc/mouse. o Percentage of lumbar BL/6 and SM/J discs at 20M, with collagen fibers in the NP. Student t-test or Mann-Whitney U test was used as appropriate except when comparing differences between groups shown in d, d’, g and o where Chi-square test was used. Data are represented as mean ± SEM. Scale bar (disc) = 200 µm.

We investigated the impact of herniations on cell phenotype, survival, and local inflammation. Unlike BL/6, SM/J mice did not show quantifiable staining of NP phenotypic marker CA3 (Fig.1h-h”) while showing an increased abundance of IL-6 in NP and AF (Fig. 1i-i”). Furthermore, SM/J presented a higher percentage of TUNEL-positive cells, indicating compromised cell survival (Fig. 1j-j”). These results show that SM/J mice experience accelerated age-dependent disc degeneration, with increased incidence of herniations accompanied by local inflammation, cell death, and loss of the NP cell phenotype.

### Aged SM/J mice evidence alterations in matrix composition and remodeling

To understand extracellular matrix (ECM) health during aging and in the context of spontaneous herniations, collagen fiber thickness and markers of disc matrix integrity (COL2, COMP, ACAN, and CS) and degradation (ARGxx and MMP13) were evaluated(*21*). Picrosirius red staining showed a lower proportion of thin fibers and a higher proportion of intermediate and thick fibers in 6M SM/J discs (Fig. 1k-k’, n-n”, Suppl. Fig.1b). While no differences were observed at 12M, at 20M a higher proportion of thin fibers, lower proportion of intermediate fibers, with significantly higher NP fibrosis was noted (Fig. 1l’-o, Suppl. Fig. 1b). Notably, collagen fiber thickness in SM/J discs remained relatively constant during aging; by contrast, in BL/6, a decrease in thin fibers and concomitant increase in intermediate and thick fibers was noted, suggesting early onset NP fibrosis in SM/J and age-associated AF collagen maturation in BL/6 (Fig. 1n-o).

COL2 showed a marked reduction in abundance in NP and AF of 20M SM/J mice (Suppl. Fig. 2a-a”); however, COMP levels were comparable between the strains (Suppl. Fig.2b-b”). Notably, abundance of ACAN, the primary disc proteoglycan, was lower in NP and AF compartments of SM/J mice (Suppl. Fig. 2c-c”); however, abundance of chondroitin sulfate, a glycosaminoglycan side chain critical to the hydration of aggrecan and other sulfated proteoglycans, was decreased in 20M SM/J NP compartment, but elevated in the AF at both 12 and 20M, a characteristic feature associated with disc degeneration (Suppl. Fig. 2f). Moreover, a higher abundance of ARGxx, a maker of aggrecan degradation, and MMP-13 was observed in SM/J discs (Suppl. Fig. 2d-e”). These results indicate that SM/J herniations coincide with an altered ECM composition and increased catabolism.

### AF tissue shows increased inflammatory changes before herniation onset

To investigate the molecular signatures associated with increased susceptibility to disc herniation, microarray analysis of the AF before the onset of herniations was performed using 12M SM/J (degenerated, non-herniated discs) and BL/6 (healthy discs) (Fig. 2a). Principal component analysis showed distinct, strain-based profiles (Fig. 2b) and 2,595 differentially expressed genes (DEGs) (46% upregulated, 54% downregulated) were identified using FDR < 0.05 and fold change (FC) ≥ 2.0 (Fig. 2c-e). To study the biological impact of these DEGs the PANTHER Overrepresentation Test was used on upregulated and downregulated DEGs. Upregulated DEGs were enriched for B and T cell activation, inflammation, apoptosis, p53, JAK/STAT signaling, and angiogenesis; and downregulated DEGs were enriched for the regulation of metabolic processes, innate response, response to stress, and ion transport (Fig. 2f, Suppl. Fig. 3 and Fig. 2F), highlighting possible contributors to AF disruption and eventual disc herniation.

**Fig. 2:**
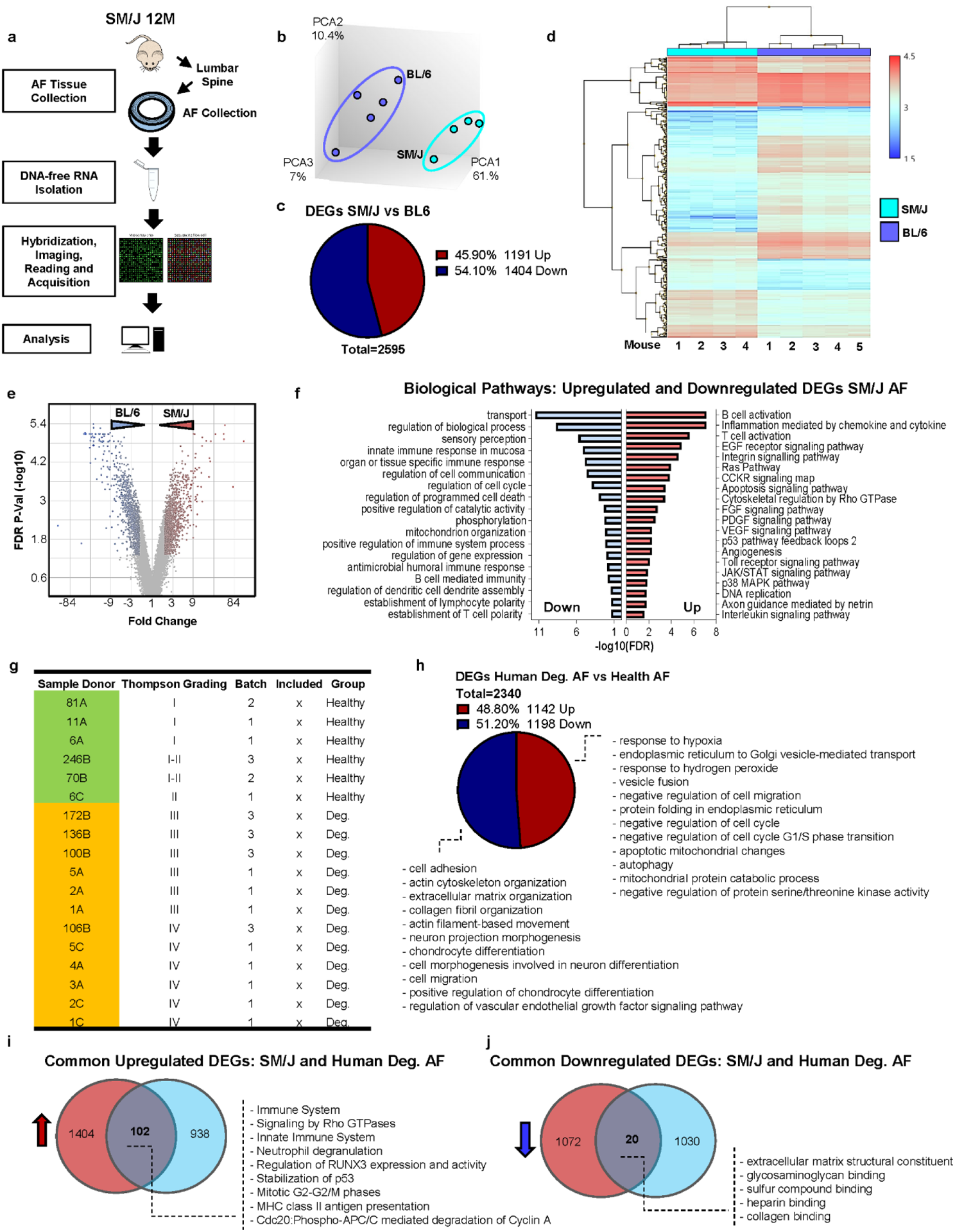
Transcriptomic signatures of human AF degeneration share similarities with SM/J mice. **(a)** Schematic summarizing study design for the transcriptomic analysis of AF tissues. **(b)** Transcriptomic profiles of BL/6 AF (n=5 mice) and SM/J AF (n=4 mice) tissues from 12M mice clustered distinctly by principal component analysis. **(c)** Pie chart showing up and downregulated DEGs between SM/J vs. BL/6, FC ≥ 2, FDR ≤ 0.05. **(d)** Hierarchical clustering of DEGs, FC ≥ 2, FDR ≤ 0.05. **(e)** Volcano Plot, showing up-and downregulated DEGs from the 12M SM/J AF vs. BL/6 AF comparison used for GO Process enrichment analysis, FC ≥ 2, FDR ≤ 0.05. **(f)** Representative GO processes from DEGs from SM/J AF vs. BL/6 AF. **(g)** Human samples from GSE70362 included in the comparative analysis with SM/J. **(h)** Representative GO processes of up and downregulated DEGs from Human Degenerated AF vs. Human Healthy AF. **(i)** Representative Venn Diagram of common upregulated DEGs and correspondent enriched GO processes from 12M SM/J AF vs. 12M BL/6 AF and Human Degenerated AF vs. Human Healthy AF. **(j)** Representative Venn Diagram of common down-regulated DEGs and corresponding enriched GO processes from 12M SM/J AF vs. 12M BL/6 AF and Human Degenerated AF vs. Human Healthy AF. GO analysis was performed using the PANTHER Overrepresentation Test with GO Ontology database annotations and a binomial statistical test FDR ≤ 0.05.

### Transcriptomic signatures of human AF degeneration share similarities with SM/J mice

To evaluate the clinical relevance of SM/J transcriptomic data, it was compared to a previously reported dataset (GSE70362) of healthy and degenerated human AF tissues (Fig. 2g)(*22*). Hierarchical clustering demonstrated distinct transcriptomic profiles between healthy and degenerated human AF samples (Fig. 2g, Suppl. Fig. 4a) Upregulated DEGs (∼49%) were associated with cell regulation of MAPK activity, responses to cytokines, hypoxia, ion transport, positive regulation of apoptotic processes, and cell cycle arrest. By contrast, downregulated DEGs (51%) were enriched for collagen fibril organization, actin cytoskeleton organization, cell adhesion, chondrocyte differentiation, and primary metabolic processes (Fig. 2h). Common DEGs between human and SM/J consisted of 102 upregulated DEGs enriched for regulation of mRNA metabolic process, negative regulation of mitotic cell cycle, and neutrophil activation involved in immune response and 20 downregulated DEGs enriched in ECM structural constituents, glycosaminoglycan binding, sulfur compound binding, and heparin-binding (Fig. 2j, Suppl. Fig. 4b, b’). These results suggest a degree of conservation between human and SM/J AF degeneration, highlighting shared molecular pathways underpinning the pathology.

### SM/J mice show altered structural integrity of subchondral bony endplate

Sclerosis, increased BMD, and decreased closed porosity of subchondral bone correlate with pain and local nerve growth in back pain and OA models(*23*). Due to this proposed relationship between the disc and the structural integrity of adjacent vertebrae and subchondral bony endplate, morphological metrics of the L3-S1 vertebral subchondral bony endplate were analyzed. 20M SM/J mice showed an accelerated osteoporotic phenotype, evidenced by decreased cortical and trabecular thickness and number(*5*). Interestingly, subchondral bony endplate volume did not differ between SM/J and BL/6 despite smaller vertebral size in SM/J (Fig. 3a-c). This finding coincided with higher tissue mineral density and cortical thickness of endplates in SM/J males and a higher bone volume fraction in females and males (Fig. 3d-f). Additionally, SM/J males showed lower bone surface complexity and closed porosity (Fig. g-h).

**Fig. 3:**
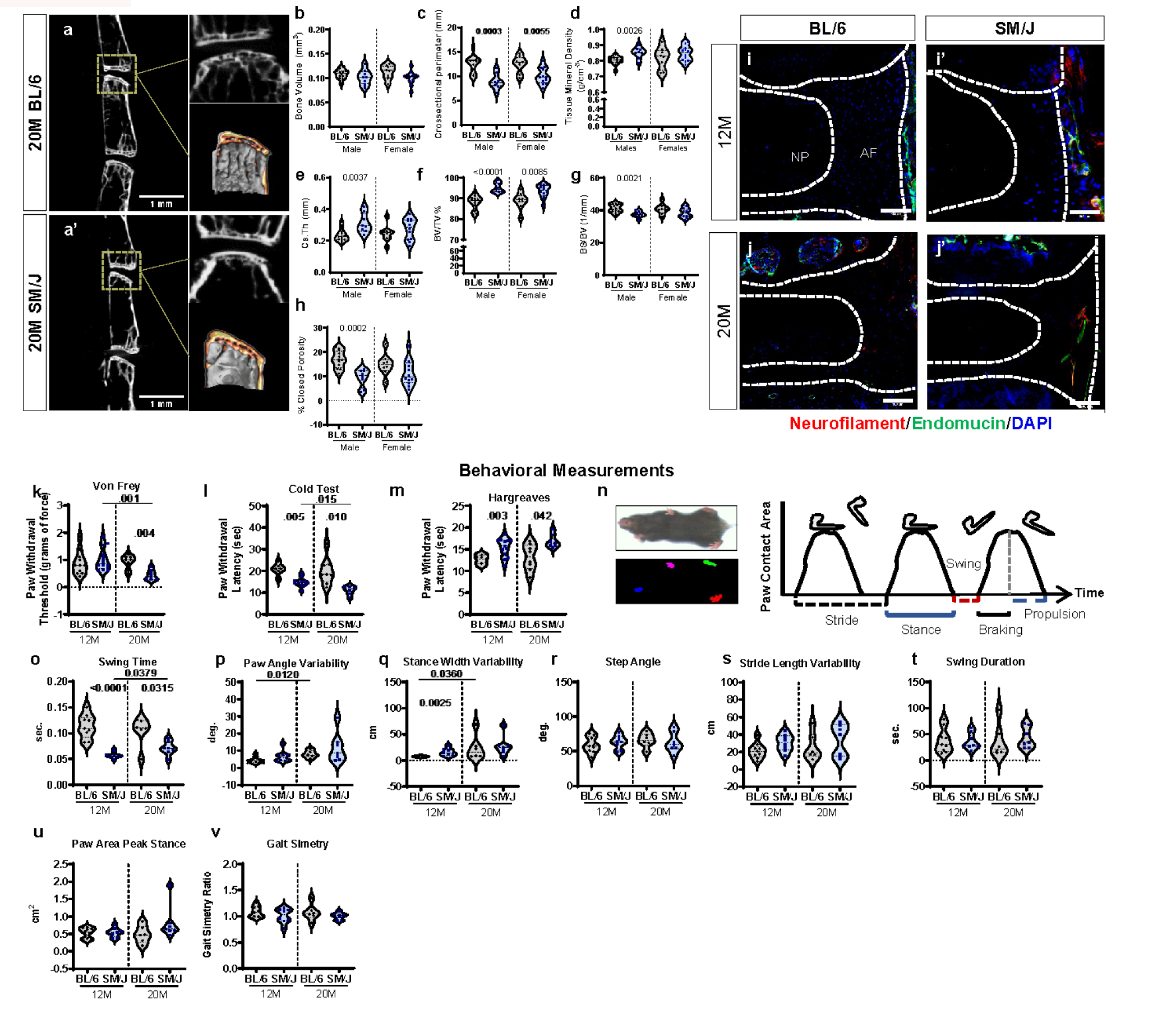
SM/J mice with aging show altered subchondral bone structure, tissue innervation, and altered sensitivity to mechanical and temperature stimuli. **(a-a’)** μCT reconstruction of 20M BL/6 and SM/J vertebrae, with the marked subchondral bone region of interest. **(b-h)** Analysis of subchondral bony endplate parameters: **(b)** bone volume, **(c)** cross-sectional perimeter, **(d)** tissue mineral density, **(e)** cross-sectional thickness, **(f)** bone volume (BV/TV), **(g)** bone surface/volume (BS/BV), and **(h)** % closed porosity between 20M BL/6 and SM/J males and females. n_BL/6female_= 2, n_BL/6male_= 3; n_SM/Jfemale_= 2, n_SM/Jmale_= 2; 4 vertebrae (L3-L6/mouse). Two-tailed t-test or Mann-Whitney test was used as appropriate. Data are represented as mean ± SEM. **(i-j’)** Characterization of neurofilament and Endomucin presence in BL/6 and SM/J mice discs at 12M and 20M. n=6 mice/strain, 2-3 levels/mouse. Scale bar = 200 µm. **(k-m)** Sensitization analysis of: **(k)** mechanical (Von Frey), **(l)** hot (Hargreaves), and **(m)** cold stimulus. n_BL/6 12M_ = 6-11, n_BL/6 20M_ = 8; n_SM/J 12M_= 11-14, n_SM/J 20M_ = 6-7. **(n)** Schematic of DigiGait gait analysis and parameters. **(o-v)** Gait parameters analyzed: **(o)** swing time, **(p)** paw angle variability, **(q)** stance width variability, **(r)** step angle, **(s)** stride length variability, **(t)** swing duration, **(u)** paw area peak stance, **(v)** gait symmetry. n_BL/6 12M_ = 9, n_BL/6 20M_ = 6; n_SM/J 12M_= 8, n_SM/J 20M_ = 7. t-test or Mann-Whitney test was used to compare differences between the groups as appropriate. Data are represented as mean ± SEM.

### SM/J mice show peripheral vascularization and innervation of the herniated discs

Disc herniation is tightly correlated with cLBP and radiculopathy. Neurofilament, a marker of axonal ingrowth(*17*), and endomucin, a marker of neovascularization commonly associated with neoinnervation(*16*), were used to investigate nerve and blood vessel ingrowth in herniated SM/J discs. We observed that endomucin and NF-stained structures were in proximity outside the disc at 12M but were present within the outer AF of herniated discs in 20M SM/J (Fig. 3i-j’). These results demonstrate an association between disc herniation and local axonal and blood vessel ingrowth, as reported in humans(*24*).

### SM/J mice show increased sensitization to mechanical and cold stimuli, along with changes in gait

Human herniation is often associated with several neuropathic symptoms(*25*). To assess similar behavioral changes, mechanical (Von Frey filament) and temperature (Hargreaves thermal; cold plantar assay) sensitization were measured in SM/J and BL/6 mice(*26*, *27*). Compared to BL/6, SM/J mice demonstrated a lower threshold for mechanical stimulation inducing a pain-like response during aging (Fig. 3k). Additionally, SM/J showed a decreased latency for a noxious cold stimulus inducing a pain-like response than BL/6 at 12M and 20M (Fig. 3l). By contrast, Hargreaves testing (heat sensitivity) showed increased desensitization in 12M SM/J mice which persisted at 20M (Fig. 3m). To study correlation of gait features with neurological pathologies or locomotion-related pain, gait analysis was performed(*28*) (Fig. 3n). SM/J mice presented an age-dependent increase in swing time; however, compared to BL/6, swing time was shorter in SM/J, independent of age. (Fig. 3o). Paw angle variability increased with age in BL/6, and a similar trend was observed in SM/J mice (Fig. 3p). Interestingly, while the stance width variability rose with age in BL/6, it remained lower than in SM/J at 12M (Fig. 3q). There were no significant changes in other parameters associated with ataxia (step angle and stride length variability), impaired mobility (swing duration), limb injury (paw area peak stance), or gait symmetry (Fig. 3r-v). These findings show that SM/J replicate clinical behaviors commonly seen in human patients with disc herniation and radiculopathy.

### SM/J DRGs show enrichment of DEGs associated with the glial cell, neuroinflammatory response, cell, matrix adhesion modulation, and immune system regulation

To study the molecular changes in DRGs concordant with herniation and behavioral changes, transcriptomic analysis of lumbar DRGs was performed. While SM/J profiles diverged with age, BL/6 profiles did not mirror this separation (Fig. 4a). Source-of-variation analysis showed the compounded effects of age and strain (“Condition”) were the highest source of variation among samples, whereas sex-based effects were negligible in both strains (Fig. 4b). Hierarchical clustering of 12M SM/J and BL/6 showed divergent transcriptomes between the strains (Suppl. Fig. 4a). There were 188 upregulated DEGs (FDR < 0.05, FC ≥ 2), which did not enrich in any biological pathways (PantherDB) (Suppl. Fig. 5b, c), whereas 167 downregulated DEGs were associated with the response to interferon-β, positive regulation of response to biotic stimulus, activation of the innate immune response, antigen processing, and presentation of peptide antigen pathways (Suppl. Fig. 5b, c). Comparison of DEGs from 20M with 12M SM/J showed distinct clustering during aging (Suppl. Fig. 6a), and a total of 401 DEGs were noted (327 upregulated, 74 downregulated) (Suppl. Fig. 6b). There was no biological enrichment within the downregulated DEGs; however, upregulated DEGs were associated with ECM organization and adhesion regulation; modulation of interleukins-1, −4, −6, −10, and −12; regulation of IFN-α and TGFβ pathway; synapse pruning; cellular response to amyloid-β; regulation of neuron death; astrocyte activation and microglial cell migration; toll-like receptor; and negative regulation of B cell receptor signaling (Suppl. Fig. 6b, c).

**Fig. 4:**
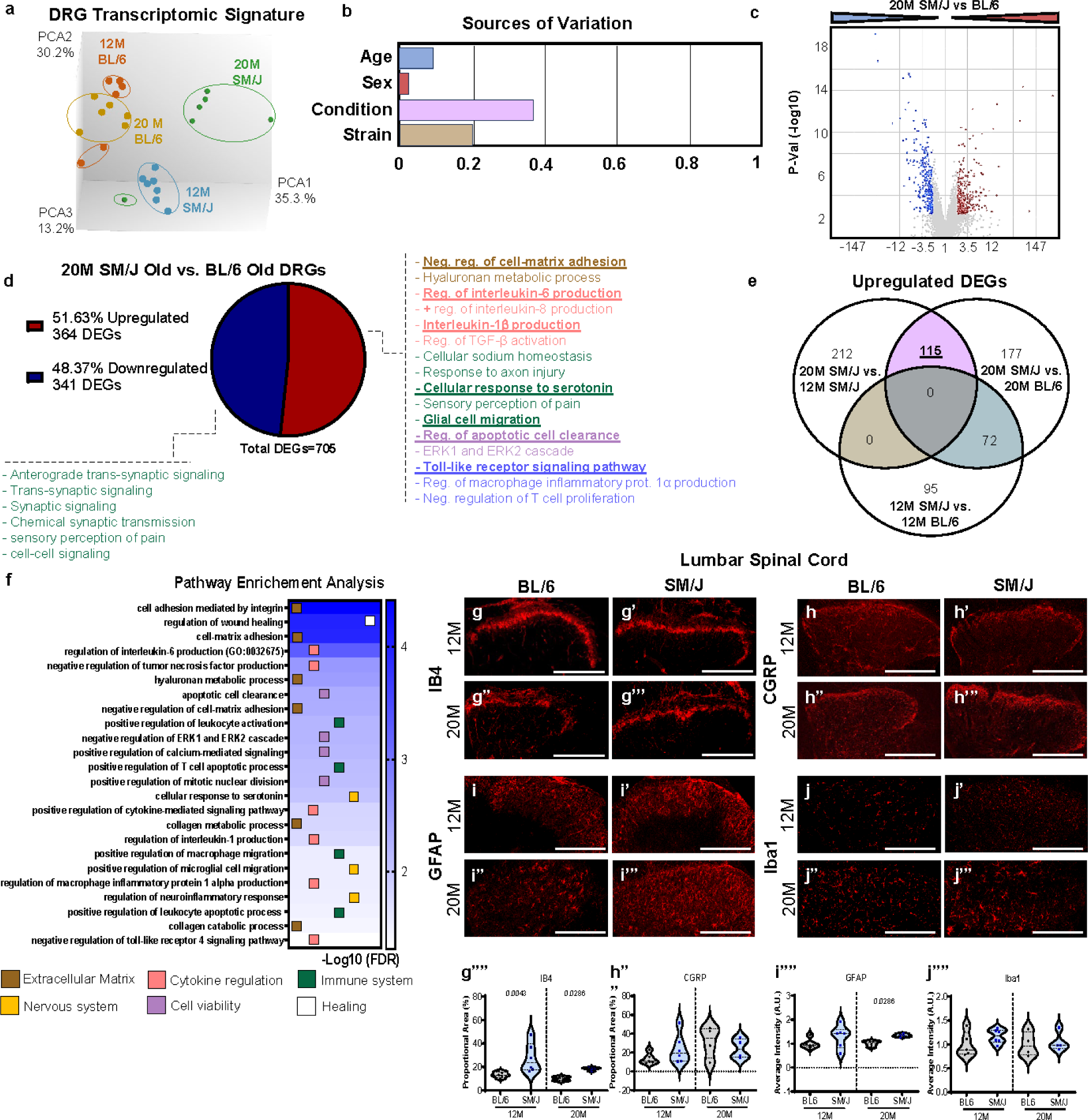
DRGs in SM/J mice show enrichment of DEGs associated with the glial cell, neuroinflammatory response, cell, matrix adhesion modulation, and immune system regulation. **(a)** Principal component analysis of transcriptomic profiles of 12M and 20M BL/6 (n=3 females, 3 males/timepoint) and SM/J (n=3 females, 3 males/timepoint) DRG tissues **(b)** weighted source of variation **(c)** volcano plot, showing up and downregulated DEGs from the 20M SM/J DRGs vs. 20M BL/6 DRGs comparison, FC ≥ 2, FDR ≤ 0.05. **(d)** Pie chart showing distribution of up and downregulated DEGs and enriched GO processes from the 20M SM/J vs. BL/6 DRGs. **(e)** Venn Diagram of common upregulated DEGs in 20M SM/J vs. 12M SM/J and 20M SM/J vs. 20M BL/6, but not present in 12M SM/J vs. 12M BL/6. **(f)** Pathway enrichment analysis of 115 DEGs identified in Fig. 4e; analysis performed in PantherDB using the statistical overrepresentation test with GO Ontology database annotations and a binomial statistical test, FDR < 0.05 **(g-j””)** quantified immunohistochemistry of the dorsal horn of spinal cord in the lumbar region **(g-g””)** IB4, **(h-h””)** CGRP, **(i-i””)** GFAP, and **(j-j””)** IBA1. n_BL/6 12M_ = 6, n_BL/6 20M_ = 4; n_SM/J 12M_= 6, n_SM/J 20M_ = 4, 2 spine levels/animal. Two-tailed t-test or Mann-Whitney test was used as appropriate. Data are represented as mean ± SEM. Scale bar A-H’ = 200 µm.

Hierarchical clustering of 20M SM/J vs. BL/6 showed clear strain-based segregation of DEGs (Suppl. Fig. 7a) with 364 upregulated and 431 downregulated DEGs in SM/J (Fig. 4d). Upregulated DEGs (Suppl. Fig. 7b) showed enrichment of negative regulation of cell-matrix adhesion and hyaluronan metabolic process; regulation of interleukins-1β, −6, −8; TGFβ activation; response to axon injury; cellular response to serotonin; glial cell migration; sensory perception of pain signaling; regulation of apoptotic cell clearance; ERK1 and ERK2 cascade; modulation of toll-like receptor signaling, macrophage inflammatory protein 1α production; and negative regulation of T cell proliferation (Fig. 4d). Downregulated DEGs (Suppl. Fig. 7b) enriched for trans-synaptic signaling, sensory perception of pain, and other signaling pathways (Fig. 4d).

To identify any DEGs associated with the pain-related behaviors exhibited by 20M SM/J and to minimize the effect of age-and strain-based variation, we investigated the DEGs common between the SM/J aging and 20M SM/J vs. BL/6 comparisons and subtracted the baseline strain-associated DEGs identified from 12M SM/J vs. BL/6 comparison (Fig. 4e and Suppl. Fig. 8). Eight downregulated transcripts: *Gp1bb, Spry3, Izumo4, Ephx4, Plpp4, Foxj1, Prrt2,* and *Arhgdig* (Suppl. Fig. 8a, b) and 115 upregulated DEGs met these criteria (Fig. 4e). Consistent with the observed pathophysiology of SM/J mice in this study, these DEGs were associated with the ECM, cytokines, nervous system, cell viability, and immune regulation pathways (Fig. 4f).

These results provide new insight into the pathological processes in DRGs in this novel model of spontaneous disc herniation.

### SM/J mice show higher staining for IB4 and GFAP in the lumbar dorsal horn

We evaluated the abundance of IB4-binding (non-peptidergic) and CGRP^+^ (peptidergic) nociceptive axons in the dorsal horn of the thoracic (adjacent to non-herniated discs) and lumbar (herniated) spinal cord(*17*). In the lumbar region, abundance of IB4-positive axons was higher in SM/J mice at 12M and 20M, whereas abundance of CGRP+ axons was maintained across all groups (Fig. 4g-h””). Microglia and astrocytes play essential roles in pain modulation, and the reactivity of these cell types in the spinal cord dorsal horn is associated with multiple chronic pain states (*18*, *29*). No differences were observed in levels of the microglial marker Iba1, but 20M SM/J showed higher levels of the reactive astrocytic marker GFAP (Fig. 4i-j”). This increase in GFAP abundance coincided with a broader spatial distribution of astrocytes in the deeper layers of the dorsal horn, something not observed in BL/6 or 12M SM/J mice. By contrast, no significant changes were observed in these markers in the thoracic region (Suppl. Fig 8d-e).

These findings suggest a possible role of IB4 nociceptive neurons and astrocytes in pain sensitization in SM/J mice with herniations.

### SM/J mice show altered plasma cytokine levels and increased T cell and neutrophil recruitment in the vertebral marrow

We assessed the impact of strain and aging on the systemic burden of inflammation. Plasma levels of IL-10, IL-15, IL-17A, IL-18A/F, IL-1β, IL-27p28/IL-30, MCP1, and MIP-3α rose during aging in BL/6 (Fig. 5a-c). While SM/J showed a similar increase in IL-17A and MIP-3α levels with aging, an increase in IL-16, IL-33, and MIP-1α was also noted (Fig. 5a-c). Interestingly, 12M SM/J had higher baseline levels of IL-21, IL-23, and IL-6 than BL/6 (Fig. 5a-c). At 20M, SM/J evidenced lower IL-2, IL-4, and IFN-γ, compared to BL/6. Levels of IL-17C, IL-22, IL-5, IL-9, IP-10, KG/GRO, and MIP2 were consistent among all groups. Notably, IFN-γ, IL-1β, IL2, and IL-4 showed a negative correlation to sensitization to mechanical stimuli (Fig. 5d). We evaluated the recruitment of B cells (CD19), T cells (CD3), neutrophils (Ly6G), macrophages (F4/80), and activated T cells and macrophages (MHCII) in vertebral marrow(*30*). While no changes were observed in CD19, F4/80, or F4/80-MHCII double-positive cells, CD3 and MHCII abundance was higher in 20M SM/J mice than in age-matched BL/6 and 12M SM/J mice (Fig. 5e-n). Higher levels of Ly6G in 20M SM/J mice relative to 20M BL6 mice were also observed (Fig. 5e-n). These studies support the idea that the immune system contributes to disc herniation-associated radiculopathy.

**Fig. 5:**
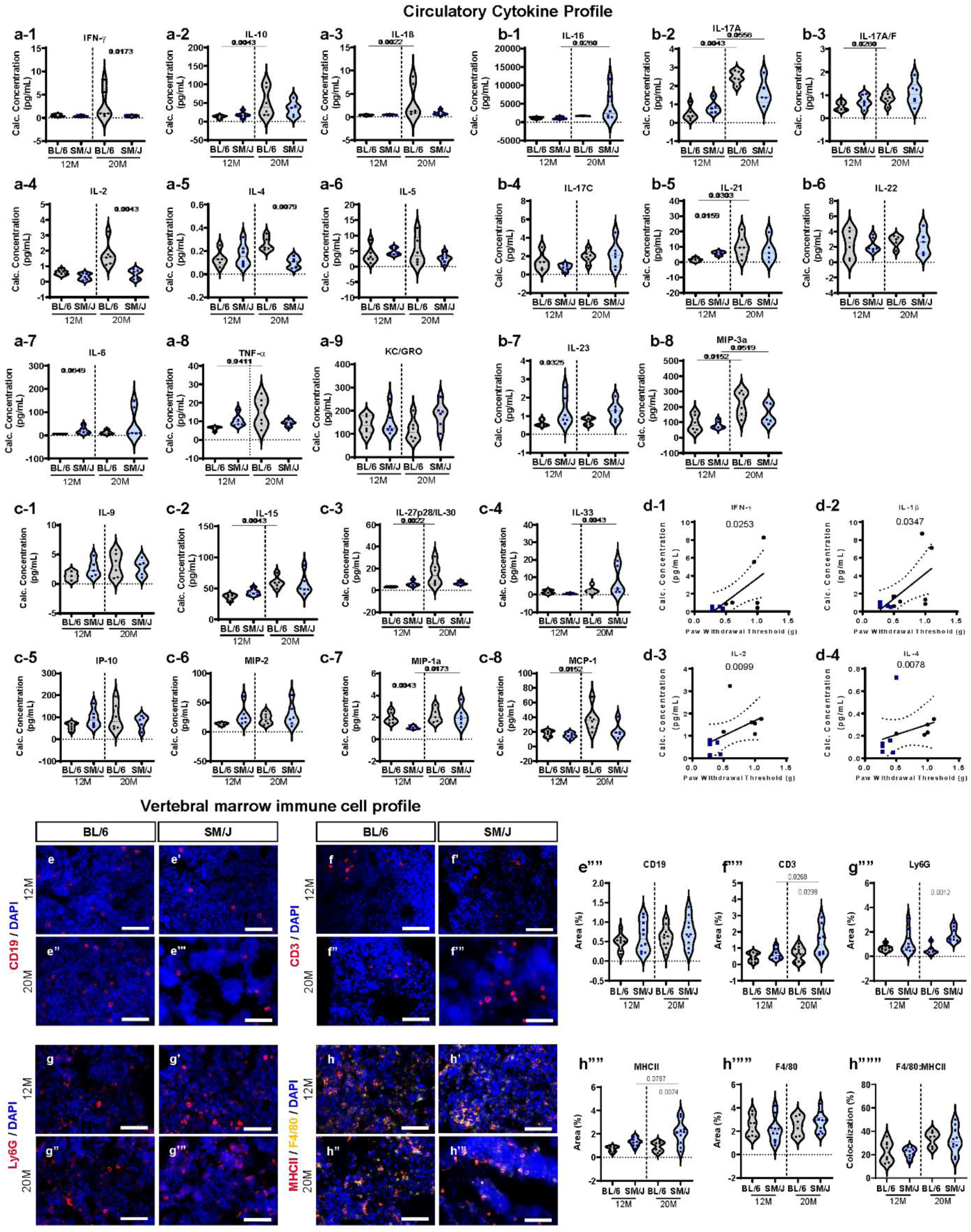
SM/J mice show altered plasma cytokine levels and increased T cell and neutrophil recruitment in the vertebral marrow. **(a1-c8)** Multiplex analysis of pro-inflammatory molecules and cytokines in plasma from 12M and 20M BL/6, and SM/J mice. t-test or Mann-Whitney test was used as appropriate. (**d1-d4)** Pearson correlation between INF-γ, IL-1, IL-2, and IL-4 plasma levels and Von Frey sensitivities. n= 4-6 mice/strain/timepoint. **(e-h”””)** quantitative immunohistochemistry of **(e-e””)** CD19, **(f-f””)** CD3, **(g-g””)** Ly6G, and **(h-h””’)** MHCII and F4/80 (individually and ratio quantification) at 12M and 20M in the vertebral bone (n= 6-10 vertebrae/strain/time point). Two-tailed t-test or Mann-Whitney test was used as appropriate. Data are represented as mean ± SEM. Scale bar A-H’ = 100 µm.

### SM/J mice show altered immune cell profiles in peripheral circulation

Immune cells are key modulators of cLBP development(*14*). To investigate the circulating immune cell profile and understand their association with pain pathology in SM/J mice, we performed a scRNA-seq of PBMCs (Fig. 6a). After normalization, the cells were grouped into 21 clusters, according to the FindNeighbors and FindClusters functions within the Seurat single-cell sequencing analysis package (Fig. 6a). Further, the data were visualized according to age and strain and cell types were labeled using the SingleR annotation tool (Fig. 6b-b’)(*31*). Dramatic changes in cell populations were observed across strains and with aging (Fig. 6c-c’, Suppl. Fig. 9a). No major changes in the distributions of cell types were observed across strains at 12 months, but by 20 months, SM/J showed lower proportions of Monocytes, and T cells, and higher proportions of B cells macrophages than BL/6 (Fig. 6c-c’).

**Fig. 6:**
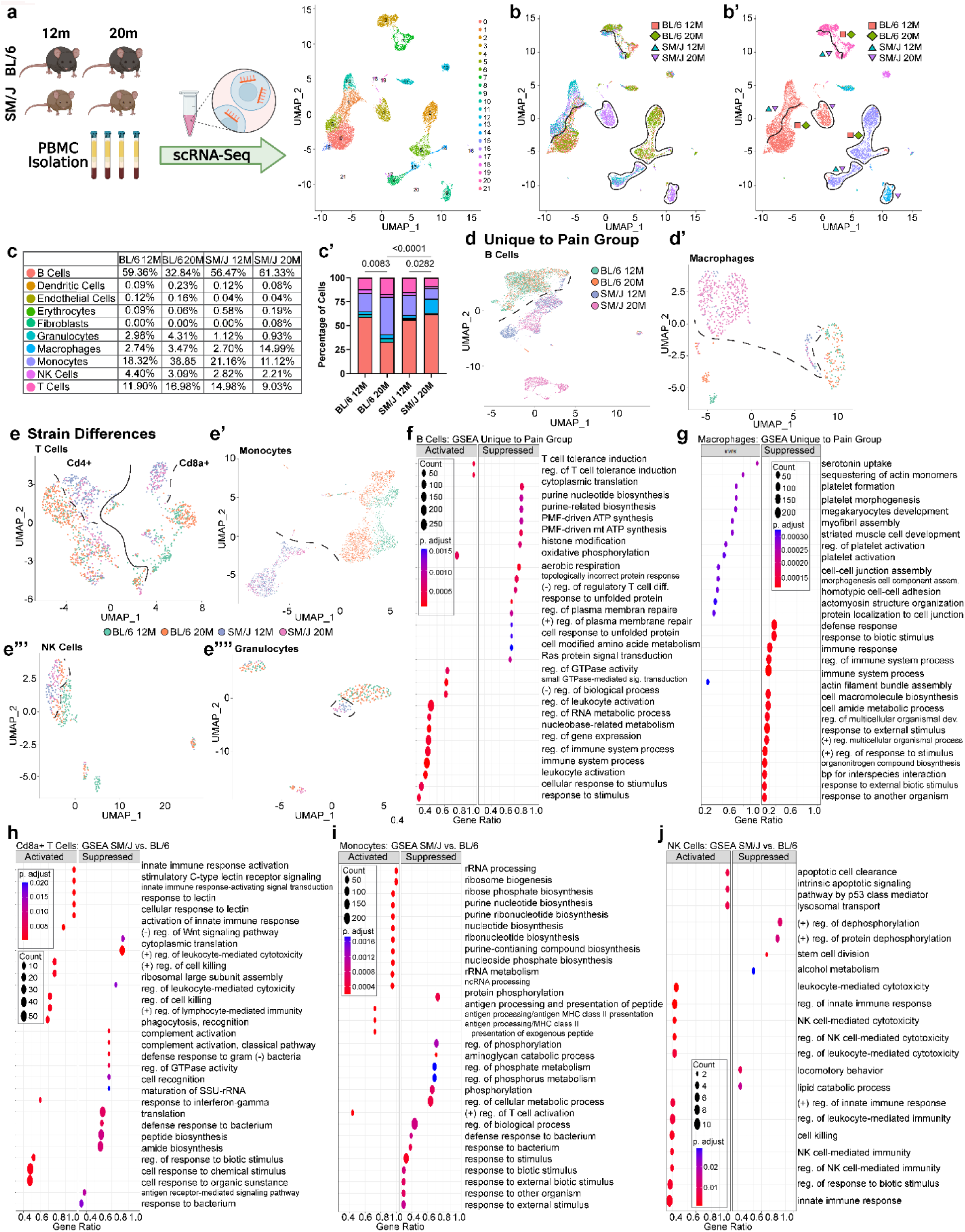
SM/J mice show altered immune cell profiles in peripheral circulation. **(a)** schematic showing experimental design and unbiased clustering of PBMC transcripts **(b-b’)** clustering of PBMC transcripts according to **(b)** strain and timepoint and after **(b’)** labeling with the SingleR program. **(c)** distribution of cell populations identified and **(c’)** table showing the proportion of each cell population. **(d-d”’)** UMAPs labeled according to mouse strain and age showing cell populations identified to be unique to SM/J at 20M, when mice show disc herniations and pain behaviors: **(d)** B Cells, **(d’)** Macrophages. UMAPs labeled according to mouse strain and age showing cell populations identified to be different between SM/J and BL/6 strains: **(e)** T Cells, **(e’)** Monocytes, **(e”)** NK cells, and **(e’”)** Granulocytes. **(e-e”)** (**f)** Gene set enrichment analysis (GSEA) in B Cells based on DEGs identified in 20M SM/J PBMCs. **(g)** GSEA of Macrophages from 20M SM/J. GSEA showing strain-based differences in **(h)** CD8a+ T cells, **(i)** Monocytes, **(j)** and NK cells. (n∼5000 cells/strain/timepoint).

Previous studies have suggested dependence of immune cell heterogeneity on inflammatory vs. quiescence state, function, and origin(*32*, *33*). To visualize heterogeneity, cell populations were subset and visualized with UMAPs, showing distinct differences in cell populations according to pain status-specific differences – B cells, and macrophages – (Fig. 6d-d’, Suppl. Fig.9b, c) and strain – T cells (subdivided into CD4+ and CD8a+ T cells), monocytes, NK cells, and granulocytes – (Fig. 6e-e’”, Suppl. Fig. 9d-g). Gene set enrichment analysis (GSEA) analysis was conducted on DEGs defined by p-value < 0.05 for each cell population using the clusterProfiler package (*34*). In B Cells of 20M SM/J, GSEA showed the activation pathways related to T cell tolerance induction, Ras protein signal transduction, and immune system processes (Fig. 6f, Suppl. Fig. 9b’-b”). On the other hand, B cells showed suppression of negative regulation of regulatory T cell differentiation and purine biosynthesis process (Fig. 6f, Suppl. Fig. 9b’-b”). Interestingly, while the proportion of macrophages in the pain group was higher, their DEGs showed suppression of defense and immune response related pathways, and activation of platelets, blood coagulation, and serotonin uptake (Fig. 6g, Suppl. Fig. 9c’-c”). T cells, despite being fewer in the 20M SM/J group, CD8+ T cells enriched for the activation of an innate response, positive regulation of leukocyte-mediated cytotoxicity and immunity, and cellular response to ifn-gamma; and for suppression of negative regulation of Wnt signaling, translation, and complement activation (Fig. 6h). SM/J CD4+ T cells showed DEGs associated with intracellular molecular regulation-related pathways (Suppl. Fig. 9d’). GSEA analysis of SM/J monocytes showed activation of positive regulation of T cell activation, antigen processing and presentation, and lymphocyte-mediated immunity; suppression of response to stimulus and regulation of biological processes (Fig. 6i). Finally SM/J NK cells enriched for the activation of the innate immune response, natural killer cell-mediated immunity, and apoptotic cell clearance (Fig. 6j).

To explore agreement between the scRNA-seq and plasma analyses, we looked for cell populations expressing *Ifng*. We found T Cells and NK cells to be the two predominantly *Ifng*-expressing populations, and trends in expression, considered with the cell population trends, confirm the reduced overall expression in 20M SM/J mice (Suppl. Fig. 10a-a”).

These analyses suggest that dysregulation of the immune system, characterized by suppression of basic immune and regulatory processes in multiple cell types, of SM/J mice contributes to chronic disc herniation and pain.

### SM/J mice show an altered immune cell profile in the spleen

To further interrogate the associations between discogenic and radicular pain and the systemic immune system modulation, cytometry by time of flight (CyTOF) analysis was conducted on splenocytes from 12M and 20M mice using 16 cell-surface markers and 8 intracellular cytokines (20M only). Using previously described immune cell signature profiles, 15 cell populations were identified based on an initial gating for CD45+ cells (Fig. 7a). This cell identification strategy also coincided with our scRNA-seq data, aligning with several populations identified in that analysis (Suppl. Fig. 10b). At 12M, SM/J showed a significantly higher frequency of T cells and NK cells than BL/6 (Fig. 7b-d). Whereas at 20M, this trend reversed for NK cells and there was no difference between two strains for overall T cell numbers. However, amongst T cells, there was a higher frequency of the CD8+ T cells in SM/J splenocytes (Fig. 7b’-d’ and Suppl. Fig. 11). Interestingly, during aging, NK cell frequency rose, and CD8+ T cell and myeloid cell frequencies decreased in BL/6 (Fig. 7b”-d”). Frequencies of NK cells, CD4+ memory T cells, and CD4+ naïve T cells decreased during aging in SM/J (Fig. 7b’”-d’”).

**Fig. 7:**
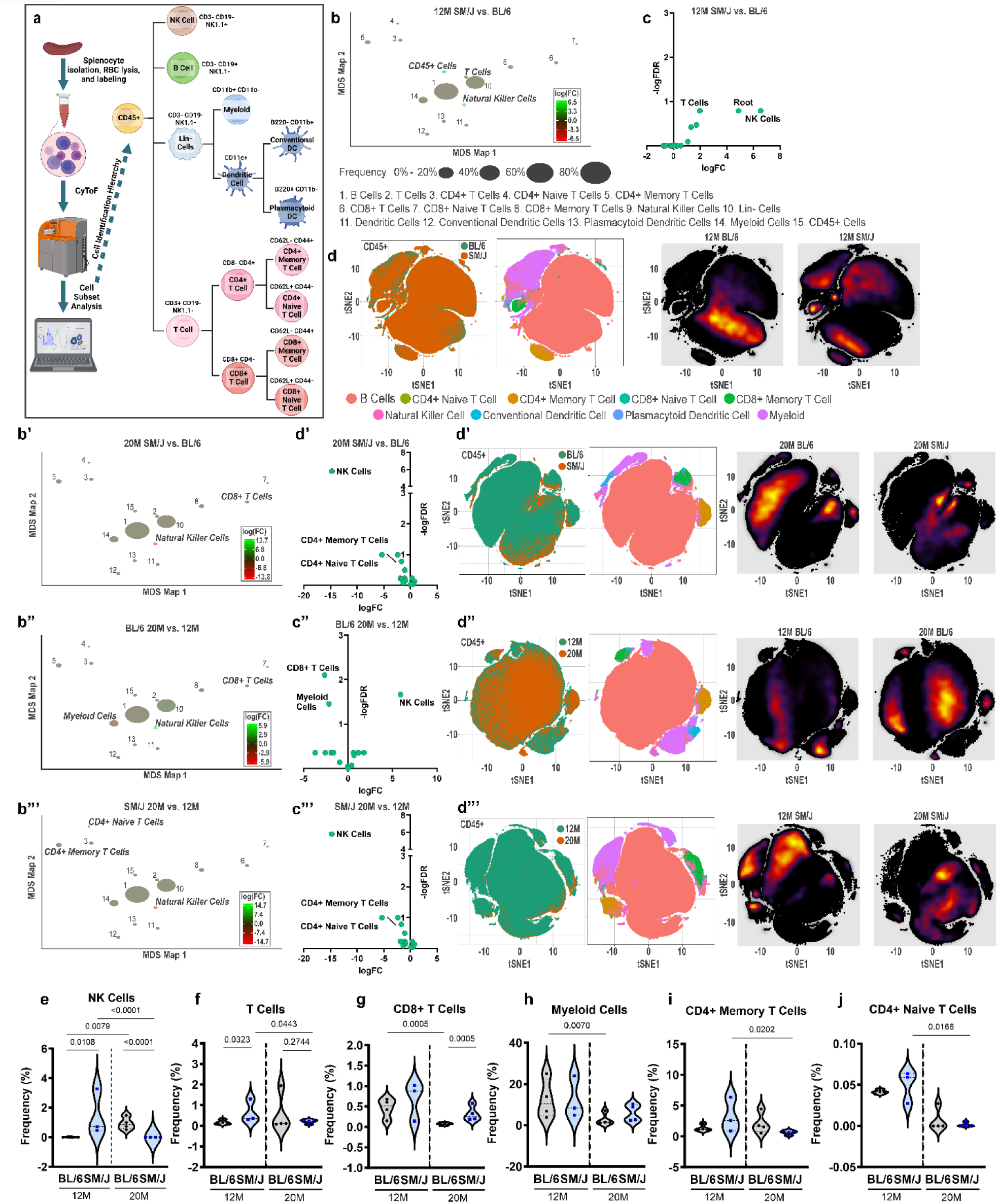
CyTOF analysis of splenocytes reveals age- and strain-based frequency differences in NK, T, CD8+ T, Myeloid, CD4+ memory T, and CD4+ naïve T cell populations. (a) schematic showing the workflow for CyTOF analysis of splenocytes from BL/6 and SM/J mice and the cell population labeling hierarchy **(b-b”’)** MSD maps showing variations in cell populations according to strain and aging **(c-c’”)** volcano plots showing cell populations in strain and age-based comparison, based on FDR and FC. **(d-d”’)** heatmaps for each comparison condition showing the distribution and relative concentration of each cell population. **(e-j)** frequency plots showing age or strain-based differences in **(e)** NK cell, **(f)** T cell, **(g)** CD8+ T cell, **(h)** Myeloid cell, **(i)** CD4+ memory T cell, and **(j)** CD4+ Naïve T cell populations. (n_12MBL/6_ = 4, n_12MSM/J_ = 3, n_20M BL/6_ = 4, n_20MSM/J_ = 4) two-tailed t-test or Mann-Whitney test was used as appropriate. Data are represented as mean ± SD.

To investigate possible functional differences in the immune cell populations in mice with herniated discs, intracellular cytokine levels were determined at 20M (Fig. 8a, Suppl. Fig. 11). At 20M, SM/J mice showed significantly higher numbers of myeloid cells producing IFN-γ (Fig. 8a, b). These inflammatory myeloid cells that include granulocytes, monocytes and macrophages (M1 macrophages) depicted that inflammatory type 1 immune response (Th1 response) characterized by IFN-γ production was predominant in SM/J mice. Lower production of anti-inflammatory Th2 cytokine IL-4 and IL-5 from in plasmacytoid dendritic cells (Fig. 8c) and CD8^+^ Naïve T cells (Fig. 8d), respectively, corroborated the inflammatory environment. IL-2 was also lower in conventional dendritic cells (Fig. 8e). Furthermore, nonspecific activation of CD8+ T cells (Fig. 8f) CD8+ memory T cells (Fig. 8g) were also observed. Overall, there was enhancement of an inflammatory Th1 response and suppression of an anti-inflammatory Th2 immune response. The Th17 immune response was also suppressed, as seen by significantly lower production of IL-17A by Lin^-^ cells (Fig. 8h) as well as myeloid cells (Fig. 8b), indicating that an inflammatory Th1 immune response, not Th17, underscored the phenotype. Significant changes in cytokine profiles of other cell populations were not observed (Suppl. Fig. 11).

**Fig. 8:**
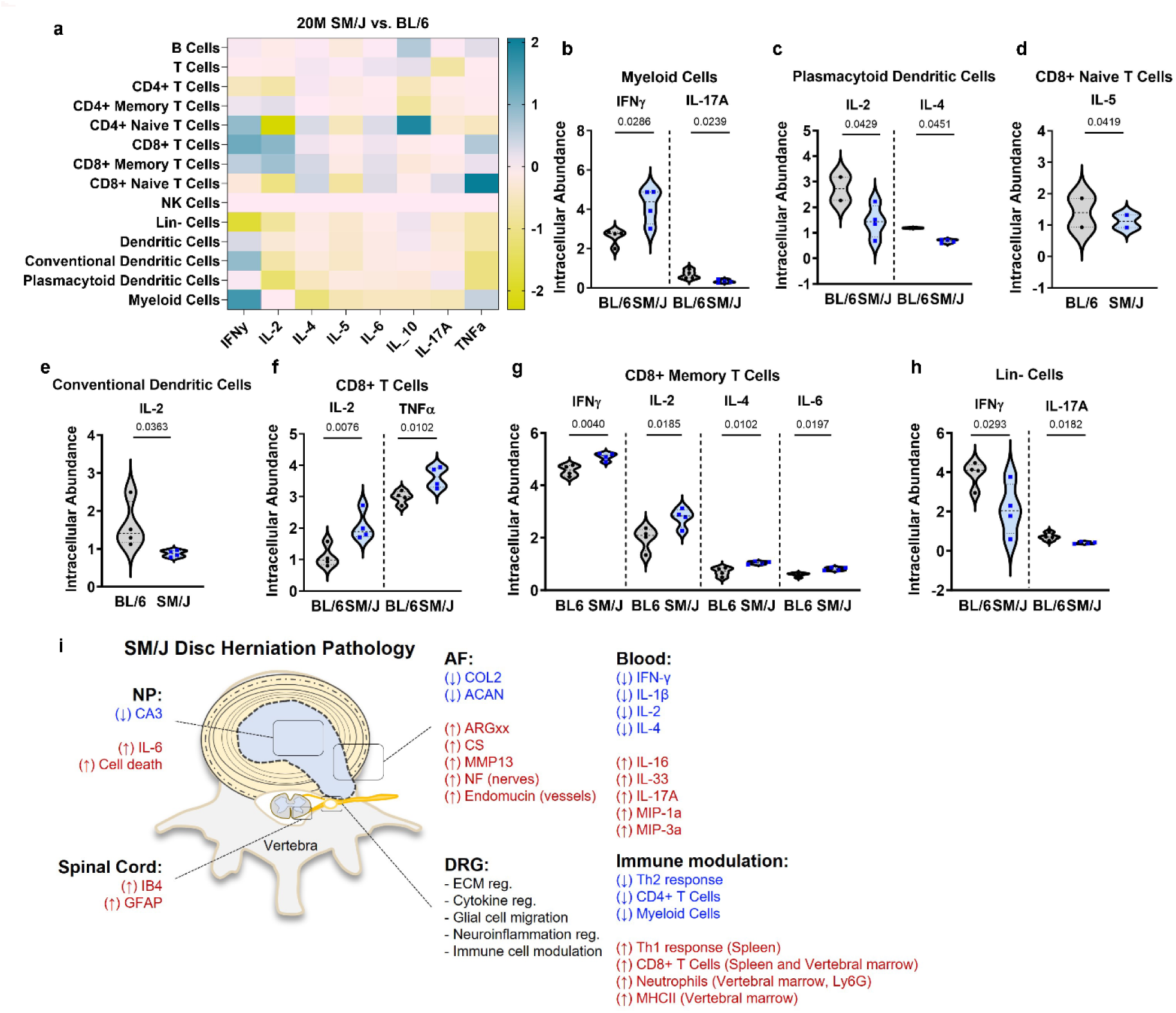
Intracellular cytokine and proinflammatory marker abundance in 20M SM/J splenocytes indicates possible repression of the innate immune system and overactivation of CD8+ T cell populations. **(a)** heatmap of splenocytes from 20M SM/J vs. BL/6 showing intracellular cytokine and proinflammatory marker abundance within each cell population identified through CyTOF analysis. (**b-h)** significant changes in cytokine/proinflammatory marker abundance in **(b)** myeloid cells, **(c)** conventional dendritic cells, **(d)** plasmacytoid dendritic cells, **(e)** Lin-cells, **(f)** CD8+ Naïve T cells, **(g)** CD8+ T cells, and **(h)** CD8+ memory T cells. (n=4 mice/strain/timepoint) two-tailed t-test or Mann-Whitney test was used as appropriate. Data are represented as mean ± SD. (i) summary schematic of phenotypic changes found in the NP, AF, DRGs, spinal cord, plasma, and immune tissues of SM/J mice relevant to herniation and chronic pain related behaviors.

## Discussion

Intervertebral disc degeneration comprises multiple degenerative phenotypes(*4*). Recently, SM/J mice were shown to evidence early onset, spontaneous disc degeneration characterized by advanced fibrosis in the caudal spine during aging, compared to BL/6 and LG/J strains that show milder fibrosis and dystrophic calcification, respectively(*5*, *20*),. These studies highlight the essential contribution of genetic background to degenerative processes, reinforcing findings from human twin studies that show familial background is a larger contributor to disc degeneration than aging and mechanical loading(*35*). This study shows that SM/J mice are highly susceptible to age-associated lumbar disc herniations. These results suggest that while genetics govern susceptibility to disc degeneration, anatomical/mechanical factors may influence and drive the final degenerative outcome(*5*). This idea aligns with studies of LG/J mice that show age-dependent calcification restricted to the caudal spine(*5*) and a significantly higher prevalence of disc degeneration at L3-5 of the human spine than the other levels (*4*).

Considering that the ECM plays a critical role in disc function, the lack of age-associated thickening of collagen fibers in SM/J suggested diminished remodeling of the AF, possibly due to low cell density (*36*). On the other hand, an increased abundance of MMP13 and ARGxx and decreased ACAN with increased CS levels suggested the breakdown of the aggrecan-rich matrix in SM/J mice, with possible compensatory efforts by other sulfated proteoglycans, such as versican(*37*). This loss of ECM integrity likely causes loss of disc structural integrity and mechanical properties(*20*). Notably, ACAN levels and turnover directly modulate axonal ingrowth and pain in degenerated human discs and arthritis models(*38*, *39*). Importantly, when AF DEGs from 12M SM/J were cross-referenced with those from humans susceptible to disc herniation, we observed a downregulation of ECM-related transcripts, further supporting a key role of ECM homeostasis in preventing herniations(*9*, *11*).

Comparing SM/J findings to those of degenerated human AF (*40*, *41*) showed dysregulation in cell cycle, cell viability, extracellular structure, and neutrophil activation pathways were shared between the two species. Previous studies have indicated the importance of the immune system in tissue healing after herniation(*42*), and while the disc compartment is immune privileged^49^, our results suggest that dysregulation of local innate immune response may precede acute herniation and prevent efficient resorption of the herniated mass and scar tissue formation. Similarly, hTNF-α transgenic mice showed higher susceptibility to disc herniation associated with compromised AF integrity and local alterations in immune cell recruitment(*11*, *43*). Notably, despite higher vertebral osteopenia in old SM/J (*5*), these mice show higher subchondral bone thickening. These results align with earlier observations, wherein an aberrant increase in subchondral bone volume/thickness in lumbar vertebrae and knee joints promoted by osteoclastic secretion of netrin-1 induces sensory innervation and pain(*15*, *23*).

cLBP is the primary clinical symptom associated with disc herniations(*44*). Additionally, patients often experience radiating pain in their limbs, increased sensitization to mechanical and thermal stimuli, and altered gait(*44*) – phenotypes we noted in aged SM/J mice. Mechanistically, degenerated SM/J discs had higher levels of IL-6, also noted in disc puncture models and osteoarthritis(*45*, *46*), which serves as a chemoattractant for axonal ingression, promoting local neuropathic pain (*47*). Accordingly, increased vascular and sensory innervations were observed in the outer AF but, unlike other studies, not in the NP of herniated SM/J discs(*48*). This finding may be explained by hernia remodeling stages: acute vs. absorbed vs. chronic(*6*, *11*). Importantly, in humans, herniated discs show presence of CGRP and IB4 positive nociceptive neurons(*24*) and an overall higher percentage of CGRP-positive peptidergic axons(*24*). Additionally, increased local IL-6 can, directly and indirectly, modulate responses in surrounding tissues, sensitizing the segmental DRGs and further perpetuating pain(*47*). In both acute and chronic pain models, ECM modulation in the DRGs correlates with pain(*49*). MMPs, including MMP2 and 9, which breakdown ECM, can interact with cytokines and integrins, reducing the local capacity for peripheral nerve regeneration, neuroplasticity, and connectivity and potentially contributing to neuropathy and chronic pain(*50*). Moreover, astrocytes interact with the ECM by expressing several matrix components, such as hyaluronan, or proliferate in response to local fibronectin singnaling(*51*, *52*). In agreement with this idea, we observed a concomitant modulation of astrocyte and matrix pathways in SM/J DRGs, with upregulated DEGs linked to microglial migration, perception of pain signaling, regulation of neuron death, and neuroinflammatory response pathways.

The spinal cord conducts pain-stimulated signals through the central nervous system to the regions of the brain (*53*). We observed a higher abundance of IB4^+^ axons, a nociceptive neuronal marker, in the dorsal horn of SM/J. In addition, we noted increased GFAP and changes in astrocyte distribution in the deeper layers of the tissue. While higher levels of GFAP are associated with increased pain, it was surprising to observe an age-associated change in the number and distribution of astrocytes in the dorsal horn in both BL/6 and SM/J strains(*29*). Expression of pain-associated markers was restricted to the lumbar region of the spinal cord, with no differences observed in the thoracic region. Coalescence of disc herniation, DRG and spinal cord changes in the lumbar spine of aged SM/J mice supports the hypothesis of a discogenic pain source in these animals.

Transcriptomic studies of the DRGs additionally highlighted the associations between discogenic sensitization, inflammatory cytokine modulation, and local immune system regulation, associations which are well-accepted(*54*). We explored the crosstalk between changes in the DRGs and systemic inflammation, as aging and senescence are associated with increased systemic inflammation, reduced immune activity and efficiency, and increased prevalence of musculoskeletal diseases and neuropathic pain(*46*, *55*, *56*). Several cytokines, such as IL-1, IL-10, IFN-γ, and TNF-α, have been associated with disc pathology(*57*), with TNF-α and IL-1 playing prominent roles in disc herniation and homeostasis(*58*). Our results showed a negative correlation between mechanical allodynia and systemic IFN-γ, IL-1, IL-2, and IL-4 levels. Considering that an insufficient and distorted immune response can limit hernia resorption and healing, these results align well with the histological evidence of unresolved herniations in aged SM/J mice. Noteworthy, local changes in several interleukins, (IL-1, −2, −4, −6, −8, −10, −12, −17) as well as IFN-γ regulation were observed in DRGs from aged SM/J mice, furthering the parallels between systemic and local inflammatory cytokines, immune regulation, and discogenic sensitization(*54*). Additionally, we found that in the AF and vertebral tissues, T cell activation correlated with persistent hernia. A previous study of herniated human discs described the presence of activated T cells in 17% of the herniated discs(*59*). Building on this framework, we analyzed immune cell profiles in the primary lymphoid organ (spleen) and peripheral circulation, which showed a contraction of the lymphoid compartment at the expense of the myeloid compartment during BL/6 aging as reported previously(*60*). Interestingly, our analyses showed an over-activation of CD8+ T cells and inhibition of CD4+ T cells in SM/J mice. These results agree with recent studies attributing a key role to CD4+ subsets, namely Treg and Th2, in suppressing pain and CD8+ T cells in chronic pain(*61*, *62*). PBMC analysis suggests a genetic background-based dysregulation of CD4+ and CD8+ cells, as SM/J mice show a unique population of these cell types by 2 year of age. Surprisingly, while we observed higher expression of INF-γ in splenic CD8+ T cells, systemic levels showed an inverse correlation with pain. While active CD8+ T cells and myeloid cells express higher INF-γ in the spleen, the systemic levels likely reflects the total decrease in INF-γ in splenic Lin-cells and PBMCs, which showed lower levels of monocytes, NK cells and T cells. Despite a lower proportion of T cells systemically, GSEA analysis suggested an elevated T cell-driven response in 20M SM/J, which aligns with findings in chronic pain patients(*62*, *63*). In line with this, SM/J splenocytes showed as immune response skewing towards Th1, supported by lower levels of IL2 and L4(*63*). Interestingly^82,8320^, sciatic nerve injury and osteoarthritis pain phenotypes are modulated by B cell recruitment and B cells expressing Mu opioid receptors (MOR) (*64*, *65*); notably SM/J mice also evidence higher proportions B cells in circulation during aging, and these cells may contribute to pain sensitization associated with herniations. As previous studies reported a vital role of innate immune system quiescence in perpetuating herniation-associated back pain, we observed a significant deficiency of innate response, specifically the monocytes, granulocytes, and NK cells in the peripheral circulation of SM/J(*14*). Importantly, while we observe an increase of peripheral macrophages in 20M SM/J group with herniations, transcriptomic profiles of showed a suppressed immune response, suggesting an insufficient immune response and possible sterile inflammation. A recent study specifically investigated disc hernia regression in the context of macrophage depletion and restoration, providing critical evidence of the need for sufficient macrophage activity in resorption(*13*). The possibility of macrophage deficiency promoting sterile inflammation was further supported by our plasma analysis, showing decreased IFN-γ and lower intracellular cytokine abundance in the splenic innate immune cell populations analyzed with CyTOF. Noteworthy, local increases of IL-6, in addition to systemic decreases of IL-2 and IL-4, are indicative of M1 macrophage polarization, which has been linked to chronic pain(*66*). Previous studies have found varying age-related trends in NK cell abundance, hypothesizing this may reflect the lack of universality in NK cell dynamics during aging(*67*). In the present study, CyTOF and PBMC analyses showed age-associated reduction of NK cells in SM/J mice, underscoring an additional indicator of dampened innate immunity. Together, these findings highlight a possible crosstalk between increased CD8+ T-cell activity and dampening of the innate immune system, driven by Th1 immune response and promoting chronic herniation by limiting hernia resolution and, consequently, the resolution of back pain.

In summary, our studies describe a novel clinically relevant mouse model of spontaneous, age-associated disc herniation on a wildtype genetic background. As in humans, genetic susceptibility plays a profound role in SM/J disc pathology and SM/J mice evidence all the primary hallmarks of discogenic chronic back pain (Fig. 8i). Importantly, our studies provide new insights into the molecular pathophysiology of chronic discogenic pain and demonstrate a key role of immune cell dysregulation in both lymphoid organs and peripheral circulation in mediating, and as a predisposing factor to, this process.

## Methods

### Mice, treatment, and study design

All animal experiments were performed under IACUC protocols approved by Thomas Jefferson University. Mice were studied at 6-(healthy adult, 6M), 12-(middle-aged, 12M), and 20 months of age (aged, 20M; mice were collected between 18 and 22 months, with an average age of 20 months). C57BL/6 mice were obtained from the rodent colony at the National Institutes of Aging (NIA), (n_BL6 6M_ = 6; n_BL6 12M_ = 15; n_BL6 20M_ = 24; n_SM/J 6M_= 6; n_SM/J 12M_= 15; n_SM/J 20M_= 22).

The SM/J mice were obtained from The Jackson Laboratory (Stock # 000687, Jackson Labs), bred, and aged at Thomas Jefferson University. The terminal point of 18-22 months was chosen based on the comparative life phases of C57BL/6J mice and humans(*68*).

### Histological Analysis

At terminal time points, mice were either euthanized with CO_2_ asphyxiation or anesthetized with ketamine followed by transcardiac perfusion with 0.9% saline, followed by 4% paraformaldehyde (PFA). Lumbar spines were dissected and fixed in 4% PFA in PBS for 6-48 hours, decalcified in 20% EDTA, and embedded in OCT or paraffin. Histological assessment was performed using 10 µm frozen or 7 µm paraffin-embedded mid-coronal sections from 6 lumbar levels (L1-S1). Spinal cord (thoracic and lumbar) and DRGs (lumbar) were harvested from PFA-perfused animals, and cryoprotected in 30% sucrose for three days, and embedded in OCT. These tissue blocks were cut serially in the sagittal planes at a thickness of 30 μm. Sections were collected on glass slides and stored at −20°C until analysis. Safranin-O/Fast Green/Hematoxylin staining was conducted on 6 lumbar levels (L1-S1) of each spine. Picrosirius red staining was used to visualize the collagen fibers within the disc. Staining was visualized using an Axio Imager 2 microscope (Carl Zeiss) using 5×/0.15 N-Achroplan or 20×/0.5 EC Plan-Neofluar objectives (Carl Zeiss) or a polarizing light microscope (Eclipse LV100 POL; Nikon, Tokyo, Japan). Four blinded observers scored mid-coronal sections from 6 lumbar discs per mouse using a modified Thompson grading scale(*20*).

### Immunohistology and cell number measurements

Following antigen retrieval, deparaffinized sections were blocked in 5% normal serum in PBS-T and incubated with antibodies against COL2 (1:400, Fitzgerald 70R-CR008), COMP (1:200, Abcam ab231977), chondroitin sulfate (1:300, Abcam ab11570); CA3 (1:150, Santa Cruz), IL-6 (1:50, Novus NB600-1131), MMP13 (1:150, Abcam ab39012). A MOM kit (Vector laboratories, BMK-2202) was used for blocking and primary antibody incubation for ARGxx (1:200, Abcam, ab3773) staining. Tissue sections were washed and incubated with species-appropriate Alexa Fluor-594 conjugated secondary antibodies (Jackson ImmunoResearch Lab, Inc.,1:700). Sections were mounted with ProLong(™) DiamondAntifade Mountant with DAPI (Fisher Scientific, P36971), visualized with Axio Imager 2 microscope using 5×/0.15 N-Achroplan or 20×/0.5 EC Plan-Neofluar objectives, and images were captured with Axiocam MRm monochromatic camera (Carl Zeiss). Staining area and cell-number quantification were performed using the ImageJ software v1.53e, last access 09/06/2020 (http://rsb.info.nih.gov/ij/) (*69*).

Slides with spinal cord sections were first warmed for 1 h at 37 °C, before three 5-minute washes in PBS to remove OCT medium. Slides were then blocked for one hour in blocking buffer (5% Donkey Serum and .1% Triton x100 in PBS), then incubated at 4°C overnight with primary antibodies against Iba1 (1:600, WAKO, AB_839504,), GFAP (1:400, Dako, Carpinteria, California; AB_10013482) diluted in blocking buffer. Slides were then washed and incubated in the appropriate secondary antibody (1:200, Donkey anti-Rabbit Alexa Fluor 647, AB_2492288 or Donkey anti-rabbit Alexa Fluor 488, AB_2313584 in blocking buffer) before being mounted with Fluorsave (Millipore Sigma) and imaged.

### Neuronal, vascular, and immune staining in the spinal column

Endomucin (1:300, Santa Cruz Biotechnology, sc-65495), Neurofilament (1:200, Biolegend, 837904), IB4 (1:200, Sigma-Aldrich L2895); CGRP (1:50, Abcam ab36001), IB4 (1:200, Sigma-Aldrich L2895); CD19 (1:50, Biolegend, 115569), CD3 (1:50, Biolegend, 100202), Ly6G (1:50; Biolegend; 127602), MHCII (1:50; Fisher Scientific; 50-112-9377), and F4/80 (1:50; Cell Signaling; 70076S) stains were imaged on a Zeiss LSM 800 equipped with a Plan Apochromat 20x, 0.8 NA air objective (Zeiss). Wavelength excitation was 488, 594, 647, or405-nm, based on antibody conjugates, and samples were captured tiled or untiled, z-stack at 1.0µm interval. Tiled images were oriented to an upright angle and cropped to 1024×1024, and brightness and contrast were linearly adjusted using ImageJ 1.52i (National Institute of Health).

### TUNEL assay

Cell viability staining was performed using the *In situ cell death detection kit* (Roche Diagnostic). In summary, sections were deparaffinized and permeabilized using Proteinase K (20 μg/mL) for 15 min, and the TUNEL assay was carried out per the manufacturer’s protocol. Sections were washed and mounted with ProLong(™) Diamond Antifade Mountant with DAPI and visualized and imaged with the Axio Imager 2 microscope.

### Digital Image Analysis

All immunohistochemical quantification was conducted in greyscale using the Fiji package of ImageJ(*69*). Images were thresholder to create binary images, and NP, AF, EP, or vertebral compartments were manually defined using the Freehand Tool. These defined regions of interest were then analyzed either using the Analyze Particles (TUNEL and cell number quantification) function or the Area Fraction measurement.

### Tissue RNA isolation and Microarray analysis

AF tissues were dissected from lumbar discs (L1-S1) of 12-month-old BL6 and SM/J animals (n_BL6_ = 5 and n_SM/J_=4). 8-10 DRGs were collected from the lumbar region (L1-S1) from 12M and 20M BL6 and SM/J mice (3 males and three females per strain/time-point). Pooled tissue from a single animal served as an individual sample. Samples were homogenized, and total DNA-free RNA was extracted using the RNeasy® Mini kit (Qiagen).

Total RNA with RIN>4 was used for microarray analysis. According to the ABI protocol, fragmented biotin-labeled cDNA was synthesized using the GeneChip WT Plus kit (Thermo Fisher). Gene chips (Mouse Clariom S) were hybridized with biotin-labeled cDNA. Arrays were washed and stained with GeneChip hybridization wash & stain kit and scanned on an Affymetrix Gene Chip Scanner 3000 7G. Quality Control of the experiment was performed in the Expression Console Software v 1.4.1. CHP files were generated by sst-rma normalization from Affymetrix.CEL files, using the Expression Console Software. Only protein-coding genes were included in the analyses. Detection above background higher than 50% was used for Significance Analysis of Microarrays (SAM), and the FDR-value was set at 5% and FC≥2. Biological process enrichment analysis was performed using the PANTHER Overrepresentation Test with GO Ontology database annotations and a binomial statistical test with FDR≤0.05. Analyses and visualizations were conducted in the Affymetrix Transcriptome Analysis Console (TAC) 4.0 software. Pathway schematic analyses were obtained using the TAC. Array data are deposited in the GEO database, GSE252383.

### Human Microarray Analysis

Human AF transcriptomic analysis was performed with the GSE70362 dataset available in the GEO database by Kazezian *et al.* (*22*). First, degenerative, and healthy clusters were obtained by hierarchical clustering of DEGs using Euclidean metrics. Human AF samples with Thompson grades I to II were grouped into a healthy category, and grades III and IV were considered degenerated. To maintain consistency between human and mouse samples, human samples with grade V (herniation) from the dataset were excluded from the analysis. DEGs, obtained with p ≤ 0.05 and FC ≥ 1.3, were used for overrepresented biological processes enrichment analysis using the PANTHER overrepresentation test, GO Ontology database annotation, and binomial statistical test with FDR ≤ 0.05.

### Plasma Cytokine Analysis

Blood was collected by intracardiac-puncture in heparin-coated syringes and centrifuged at 1500 rcf at 4°C for 15 min to isolate the plasma. Samples were stored at −80°C until the time of analysis. According to the manufacturer’s specifications, levels of pro-inflammatory proteins, cytokines, and Th17 mediators were analyzed using V-PLEX Mouse Cytokine 29-Plex Kit (Meso Scale Diagnostics, Rockville, MD).

### Behavior testing

Modified versions of the Hargreaves, cold stimulus and von Frey filament tests were used to measure heat, cold and mechanical sensitivity, respectively(*26*, *27*). Hind paw withdrawal for Hargreaves, cold test, and von Frey was defined by removing the paw from the infrared, dry ice (through a glass plate), or mechanical stimulus. All scored withdrawal responses were associated with one or more supraspinal responses, including vocalization and licking and/or guarding of the tested paw. Spontaneous movements were not recorded, and each trial was conducted in triplicate. Cut-offs of 30s for the Hargreaves and cold stimuli tests and 2g of force for the von Frey test were used to prevent damage to the animal’s paw.

### Subchondral bone microCT analysis

Lumbar spines were dissected from 20M C57BL/6J and SM/J mice (n_BL6_= 5 and n_SM/J_= 5), fixed in 4% PFA for 48 hours and rehydrated with 1X PBS for assessment with the SkyScan 1275 μCT imaging system (Bruker). Scanning settings were 200uA current, 50kV voltage, and15μm per pixel. The skeletal parameters assessed by μCT followed previously published nomenclature guidelines (Bouxsein et al., 2010). Images were reconstructed through NRecon (Bruker) prior to analysis in CTAn (Bruker). The regions of interest were limited to subchondral bone and endplate by selecting 5 cross-sectional slices which excluded extraneous trabeculae of the vertebral body. Density of mineralized tissue (TMD) was obtained through CTAn using appropriate thresholding. Subsequent 2D analysis was used to assess bone volume (BV), mean bone area of selected cross-sections (B.Ar), cross-sectional thickness (Cs.Th), bone surface fraction (BS/BV), closed porosity(*70*).

### PBMC single-cell preparation and sequencing

Whole blood was collected from 12M and 20M SM/J and BL6 mice via intracardiac puncture and diluted into an equal volume of Dulbecco’s PBS + 2% FBS (StemCell Technologies, 07905). According to the manufacturer’s specifications, blood was layered on top of Lymphoprep™ (StemCell Technologies, 07851) in a 15 mL tube and centrifuged at 800 rcf for 20 minutes at room temperature. The mononuclear cells were washed twice with PBS + 2% FBS, and platelets were removed with an additional centrifugation at 120 rcf for 10 minutes. To facilitate removal of any remaining RBCs, a lysis step was performed (Biolegend, 420302). Briefly, lysis buffer (1X) was added to the cell suspension and incubated for 5 minutes, with periodic inversion. Cells were then centrifuged at 350 rcf for 10 minutes, and PBMCs were washed twice with PBS + 2% FBS. Cell suspensions were processed for single-cell RNA-sequencing using the 10X Genomics 3’ v3 kit (10x Genomics, Pleasanton, CA), with inter-strain multiplexing, according to manufacturer specifications. Sequencing was performed with Illumina NextSeq 2000 according to the manufacturer’s instructions (Illumina).

### scRNA-seq data analysis

Raw data were aligned to the mouse reference transcriptome (mm10-2020-A) and underwent preprocessing with Cell Ranger Multi v7.0.1. According to the authors’ recommended workflow, the count matrix was analyzed using Seurat v4.3.0)(*71*). Briefly, data were converted into Seurat Objects using the CreateSeuratObject function, and all cells were analyzed for their unique molecular identifier (UMI) and mitochondrial gene fractions; cells were included if 200 < n_feat_ < 2500, and mitochondrial genes accounted for < 5%. Data were integrated with logarithmic normalization (LogNormalize) and organized according to variation in the data using FindVariableFeatures. Dimensional reduction was performed using principal component analysis and, and the FindNeighbors (dims = 1:15) function, followed by the FindClusters functions were used to calculate the number of cell clusters using the Louvain algorithm, with the resolution parameter set to 0.5. DimPlot was then used to visualize the data as uniform manifold approximation and projections (UMAP). Cell types for each age and strain condition were determined using SingleR, with the *MouseRNAseqData* reference dataset and *main* labels(*31*). Individual cell types were individually subset and analyzed via the same methodology to determine strain and age-based variation within each cell type. The clusterProfiler package was used to conduct gene set enrichmenst analysis on DEGs (FDR < 0.05) identified in various strain and age-based comparisons(*34*). Data are deposited and may be accessed in the GEO database, GSE252474, and relevant R scripts can be found at https://github.com/mrisbudlab.

### Splenocyte preparation and CyToF

Spleens were collected from 12M and 20M SM/J and BL6 mice (n_12MBL6_ = 4, n_12MSM/J_ = 3, n_20M BL6_ = 4, n_20MSM/J_ = 4). Splenocytes were dissociated from the tissue matrix by gentle compression through a 100 μm strainer (StemCell Technologies, 27217). RBC lysis (Biolegend, 420302) was performed, followed by two PBS + 3% FBS washes. Cells were then resuspended in complete medium (Sigma Aldrich, D6046) + 10% FBS (Sigma Aldrich, F2442) + 2% 50x penicillin/streptomycin (Corning, 30-001-CI) at a concentration of 1 x 10^6^ cells/mL. To stimulate intracellular cytokine production, 2μL/mL of Cell Activation Cocktail with Brefeldin A (BioLegend, 423303) was incubated with the cells at 37°C for 4 hours. After incubation mixture were washed twice in serum free media and resuspended in serum-free media at a concentration of 2 x 10^7^ cells/mL. To label dead cells, Cell-ID™ Cisplatin (Standard Biotools, 201064) was added to each cell suspension at a concentration of 10μM. Cell staining according to the manufacturer’s protocol for the Maxpar® Mouse Sp/LN Phenotyping Panel Kit (16 markers: Ly6G/C[Gr1], CD11c, CD69, CD45, CD11B[MAC1], CD19, CD25, CD3e, TER-110, CD62L, CD8a, TCRI^2^,

NK1.1, CD44, CD4, and B220) (Standard Biotools, 201306) was carried out for all samples. 18M SM/J and BL6 splenocytes underwent a subsequent round of staining according to manufacturer’s specs for the Maxpar® Mouse Intracellular Cytokine I Panel Kit (8 markers: IFNγ, IL-2, IL-4, IL-5, IL-6, IL-10, IL-17A, TNFα) (Standard Biotools, 201310). After all staining, samples were washed, fixed with 1.6% paraformaldehyde in PBS, and incubated with Cell-ID Intercalator-Ir staining, per the Maxpar staining protocols. After this incubation, cells were washed once, passed through a 100um cell strainer, and resuspended in freeze media comprised of 90% FBS + 10% DMSO. Cells were stored at −80 ^0^C until the time of analysis. Cells were quick thawed at 37 ^0^C and washed 3 times at 800 g for 10 mins, in CAS buffer. Cells were re-suspended at 1×10^6^ per ml in CAS buffer and strained through a 35 µm filter prior to acquisition. Each sample was acquired on a CyTOF 2 Helios mass cytometer.

### CyToF Analysis

Single-cell data has been clustered using the FlowSOM R package and labeled using the Ek’Balam algorithm(*72*, *73*). Cell subset definitions follow guidelines by Maecker et al.(*74*), and Finak et al.(*75*). Cluster labeling, method implementation, and visualization were conducted through the Astrolabe Cytometry Platform (Astrolabe Diagnostics, Inc.). The Astrolabe Mass Cytometry Platform is a cloud-based service for analyzing mass cytometry data. Astrolabe labels events into immune subsets in a two-stage process. First, events are clustered using self-organizing maps implemented by the FlowSOM package. Clusters are then labeled, following a manually curated gating hierarchy. MDS maps were generated using the cmdscale R function(*76*). Differential abundance analysis was done using the edgeR package(*77*) following the methods outlined by Lun et al.(*78*). Differential expression analysis was done using the limma R package(*79*) followed by diffcyt(*80*).

### Statistical analyses

All statistical analyses were performed using Prism7 (GraphPad, La Jolla).Data distribution was assessed with the Shapiro-Wilk normality test, and the differences between the two groups were analyzed by t-test or Mann-Whitney, as appropriate. Differences between more than two groups were analyzed by ANOVA or Kruskal-Wallis for non-normally distributed data, followed by a Dunn’s multiple comparison test. χ^2^ test was used to analyze the differences between the distribution of percentages. p≤0.05 was considered a statistically significant difference.

## Acknowledgments

We thank Anshul Kumar for his assistance with tissue processing.

## Author contributions

Research design: EJN, OKO, EB, ASD, MSD, ACL, MVR

Data collection and analysis: EJN, OKO, EB, VM, ASD, MDS, VAT, AM

Manuscript preparation: EJN, OKO, MVR, ASD, ACL

## Data and materials availability

The microarray and scRNA-seq datasets that supports the findings of this study is openly available in the GEO database at GSE252383 and GSE252474, respectively. Relevant R scripts can be found at https://github.com/mrisbudlab.

