## Supplementary Figures for "Neuroimmune changes underscore pain-associated behaviors and disc herniations in SM/J mice"

Supplementary Figure 1

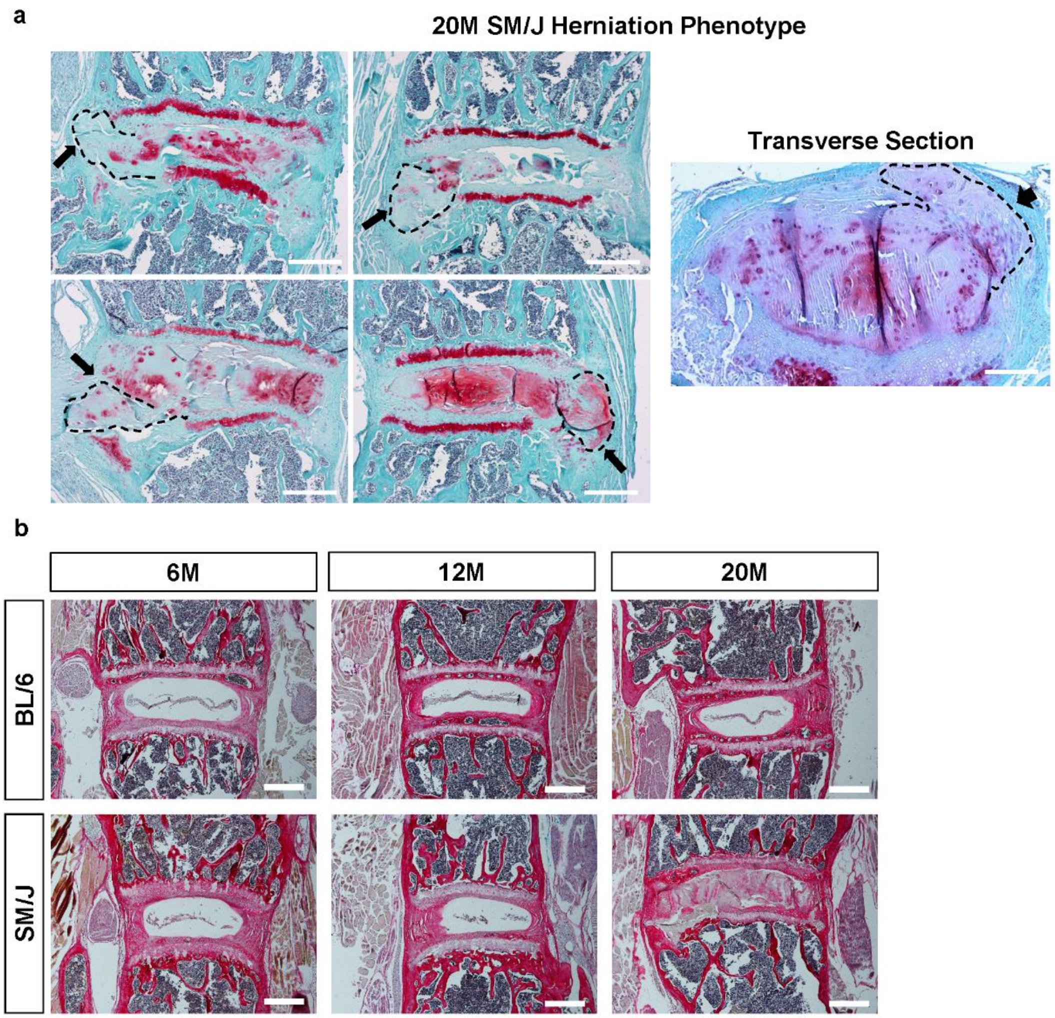


**Supplementary Figure 1:**

**(a)** Representative images showing disc herniation phenotype in 20M SM/J mice in coronal and transverse sections. **(b)** Picrosirius Red staining imaged in the brightfield showing robust staining of NP compartment in 20M SM/J mice indicative of tissue fibrosis. Scale bar = 200 µm.

Supplementary Figure 2

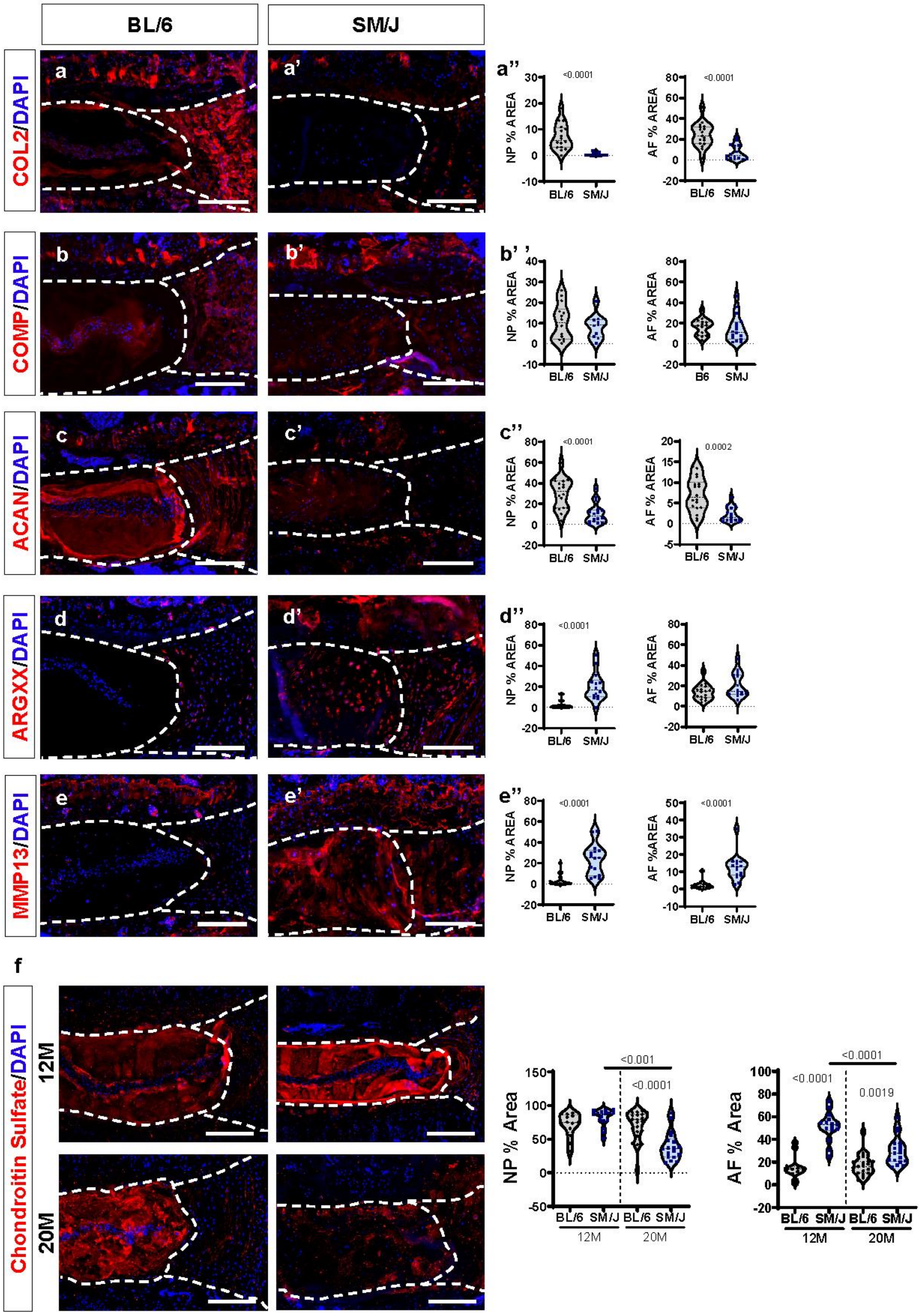

**Supplementary Figure 2:**

(**a-e")** Quantitative immunohistochemistry of 20M SM/J and BL/6 mice for **(a-a")** COL II (collagen II), **(b-b")** COMP (cartilage oligomeric matrix protein), **(c-c")** ACAN (aggrecan), **(d-d")** ARGxx (ADAMTS cleaved-aggrecan neoepitope), and **(e-e")** MMP13 (matrix metalloproteinase 13). (f) quantitative immunohistochemistry for CS (chondroitin sulfate) of 12M and 20M SM/J and BL/6 discs. t-test or Mann-Whitney test was used for 2 group comparison as appropriate. For CS, ANOVA or Kruskal-Wallis test was used as appropriate. BL/6: 12M (n=5-6 mice), 20M (n=5-6). SM/J: 12M (n=5-6), 20M (n=5-6), 2-3 disc levels/animal. Data are represented as mean ± SEM. Scale bar = 200 μm.

Supplementary Figure 3

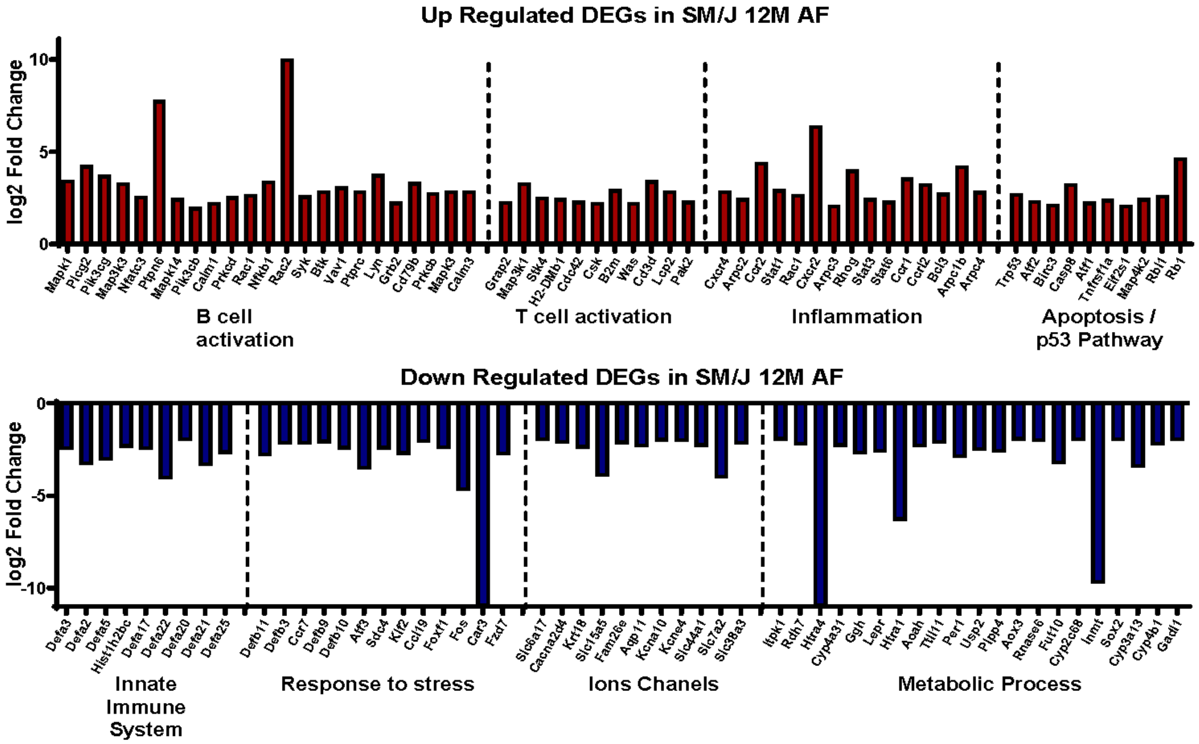


**Supplementary Figure 3:**

Representative up and downregulated DEGs in 12M SM/J AF relative to BL/6, organized according to GO processes. BL/6 AF (n=5 mice) and SM/J AF (n=4 mice).

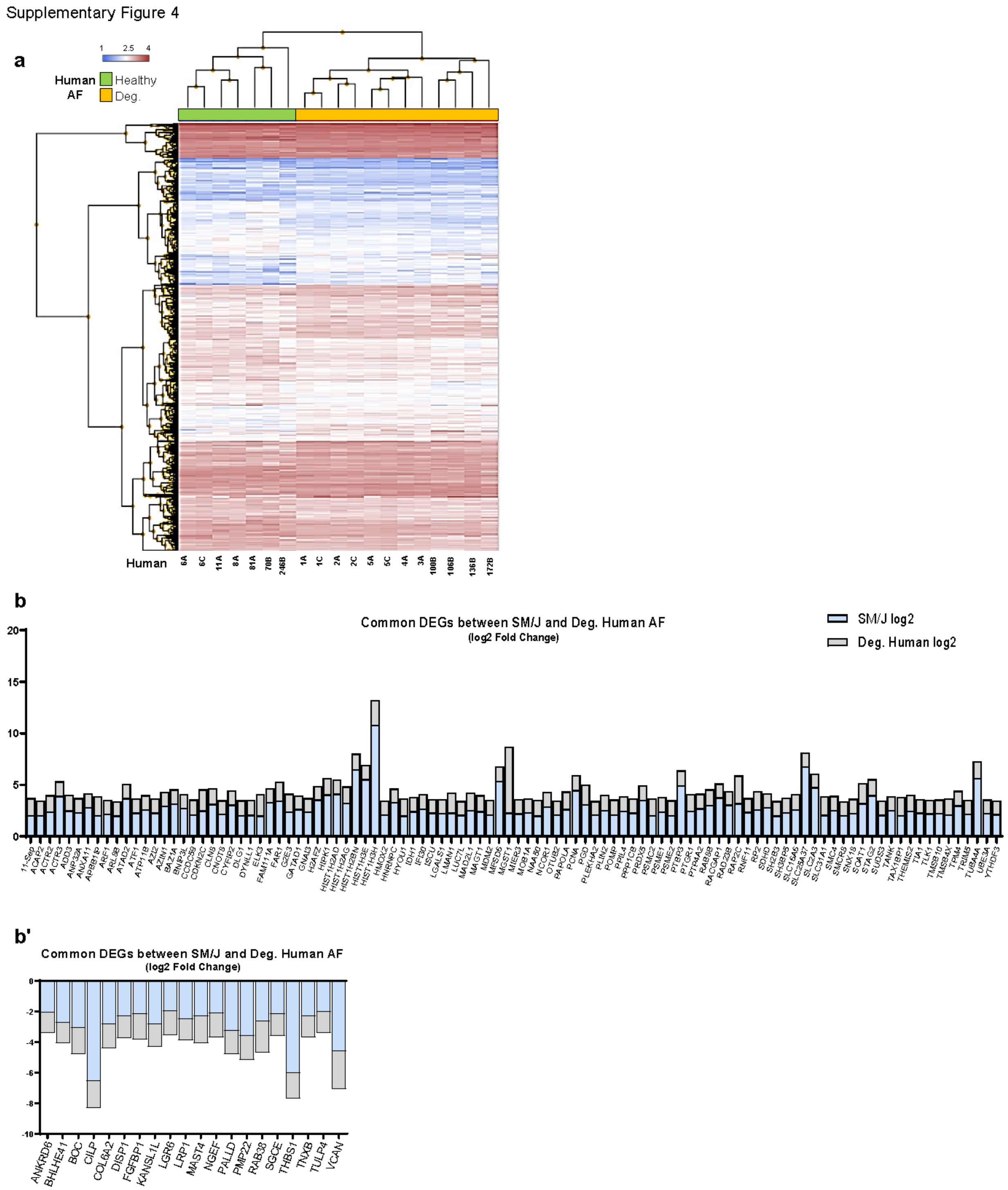


**Supplementary Figure 4:**

**(a)** Hierarchical clustering of DEGs from Degenerated Human AF vs. Healthy Human AF, FC $\geq$ 1.3, p-value $\leq$ 0.05. **(b)** Downregulated and (b') Upregulated DEGs shared by 12M SM/J AF and degenerated human AF.

Supplementary Figure 5

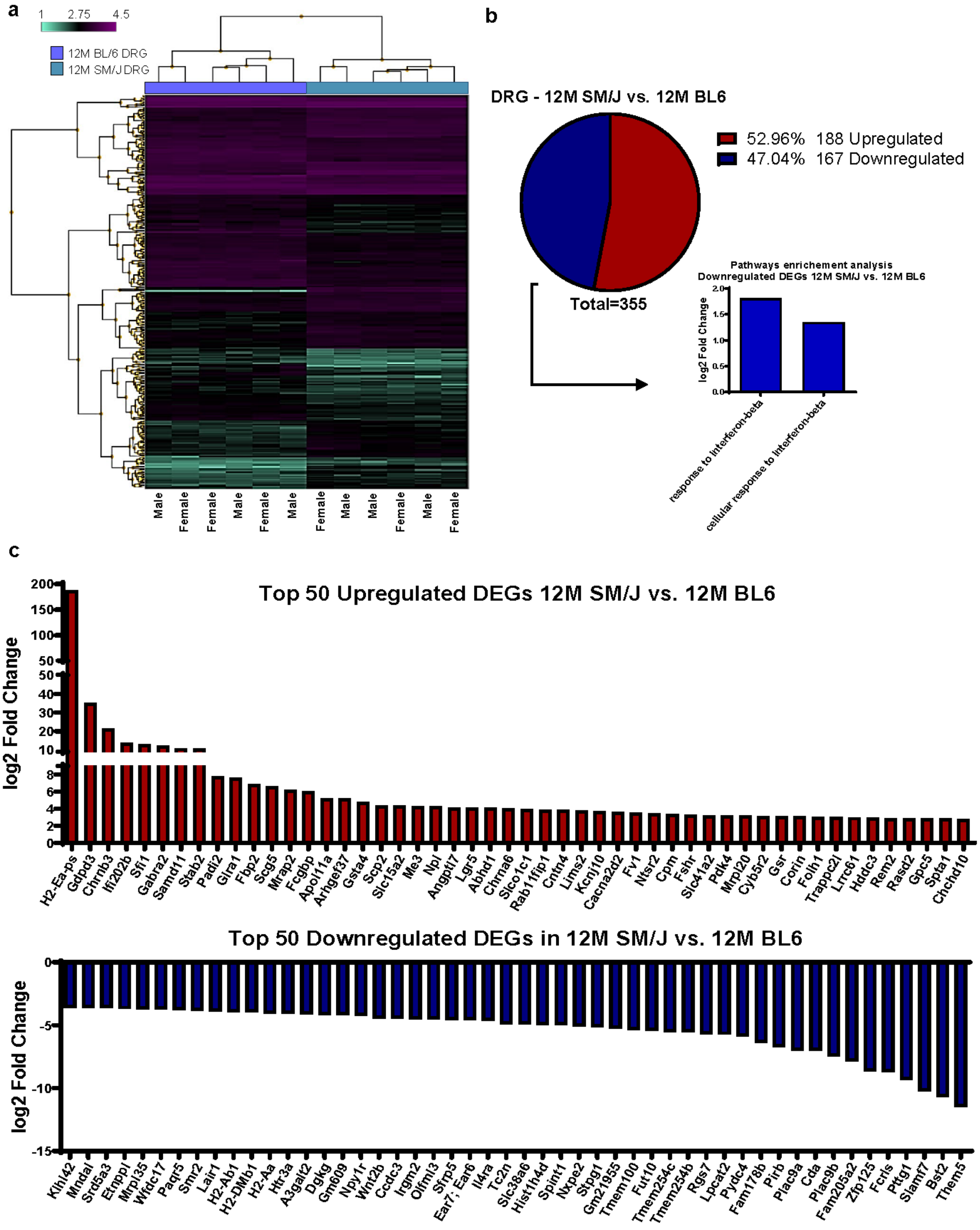

**Supplementary Figure 5:**

**(a)** Hierarchical clustering of DEGs from 12M SM/J vs. 12M BL/6 DRGs. **(b)** Pie chart showing distribution of up and downregulated DEGs and corresponding GO processes, FC $\geq$ 2, FDR $\leq$ 0.05. **(C)** Top 50 up and downregulated DEGs from 12M SM/J vs BL/6 DRGs. (n=3 female, 3 male mice/strain) GO analysis was performed using the PANTHER Overrepresentation Test with GO Ontology database annotations and a binomial statistical test FDR $\leq$ 0.05.

Supplementary Figure 6

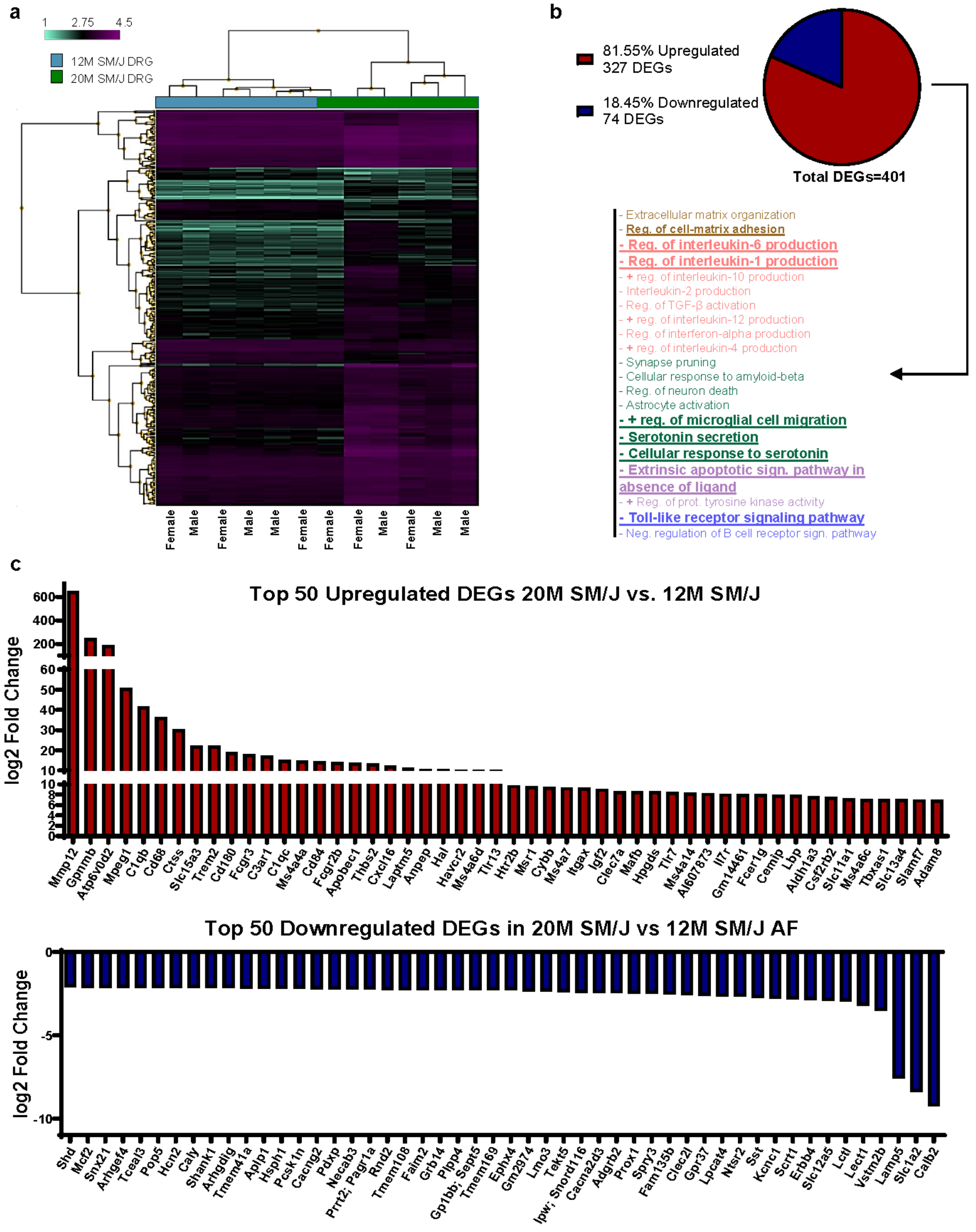

**Supplementary Figure 6:**

**(a)** Hierarchical clustering of DEGs from 20M SM/J vs. 12M SM/J dorsal root ganglia (DRGs). **(b)** Pie chart showing distribution of up and downregulated DEGs and corresponding GO processes, FC $\geq$ 2, FDR $\leq$ 0.05. **(C)** Top 50 up and downregulated DEGs from 20M SM/J vs 12M SM/J DRGs. (n=3 female, 3 male mice/timepoint) GO analysis was performed using the PANTHER Overrepresentation Test with GO Ontology database annotations and a binomial statistical test FDR $\leq$ 0.05.

Supplementary Figure 7

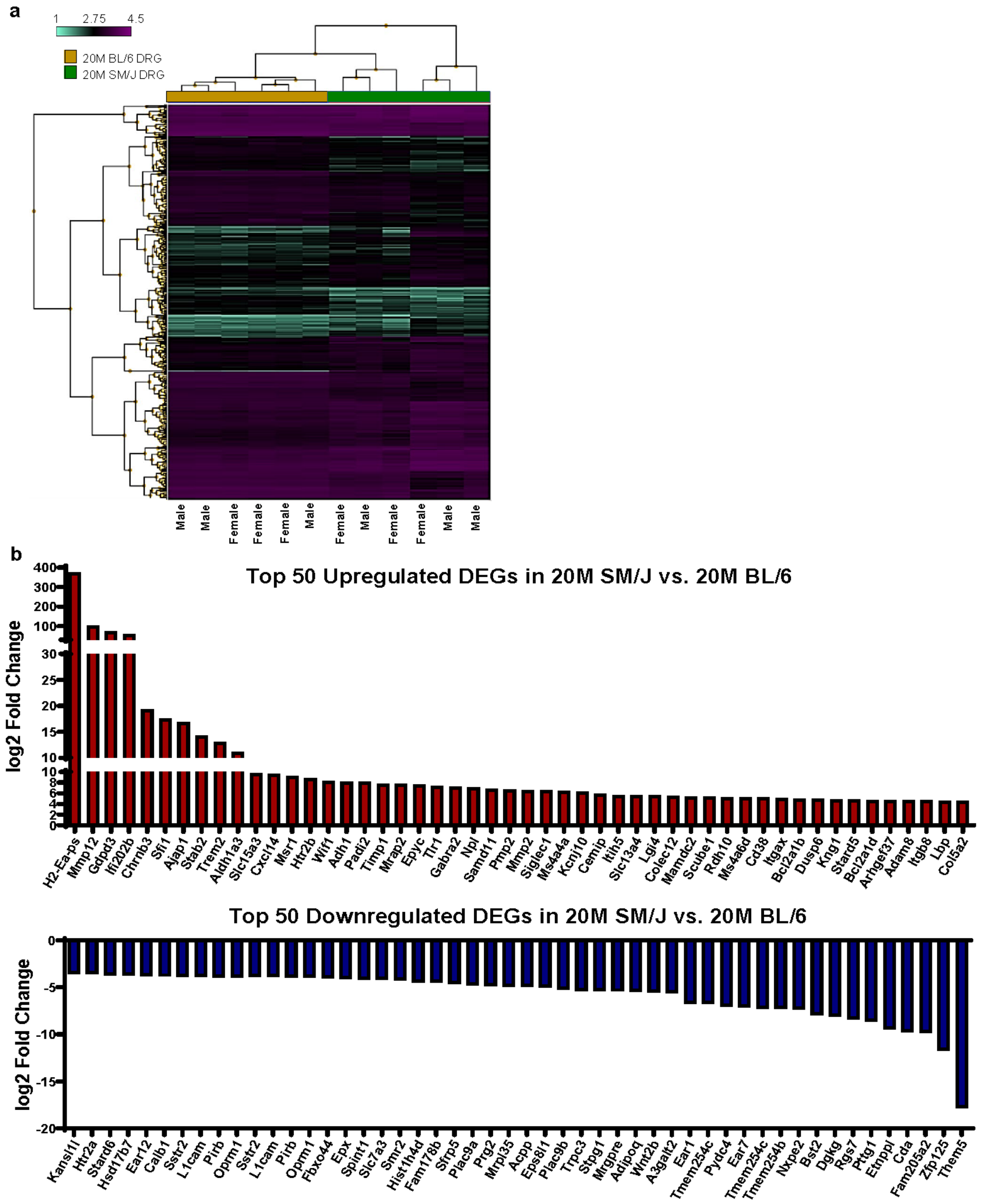

**Supplementary Figure 7:**

**(a)** Hierarchical clustering of DEGs from 20M SM/J vs. 20M BL/6 DRGs. **(b)** Pie chart of up and downregulated DEGs and corresponding enriched GO processes, FC $\geq$ 2, FDR $\leq$ 0.05. **(C)** Top 50 up and downregulated DEGs from 20M SM/J vs BL/6 DRGs. (n=3 female, 3 male mice/strain) GO analysis was performed using the PANTHER Overrepresentation Test with GO Ontology database annotations and a binomial statistical test FDR $\leq$ 0.05.

Supplementary Figure 8

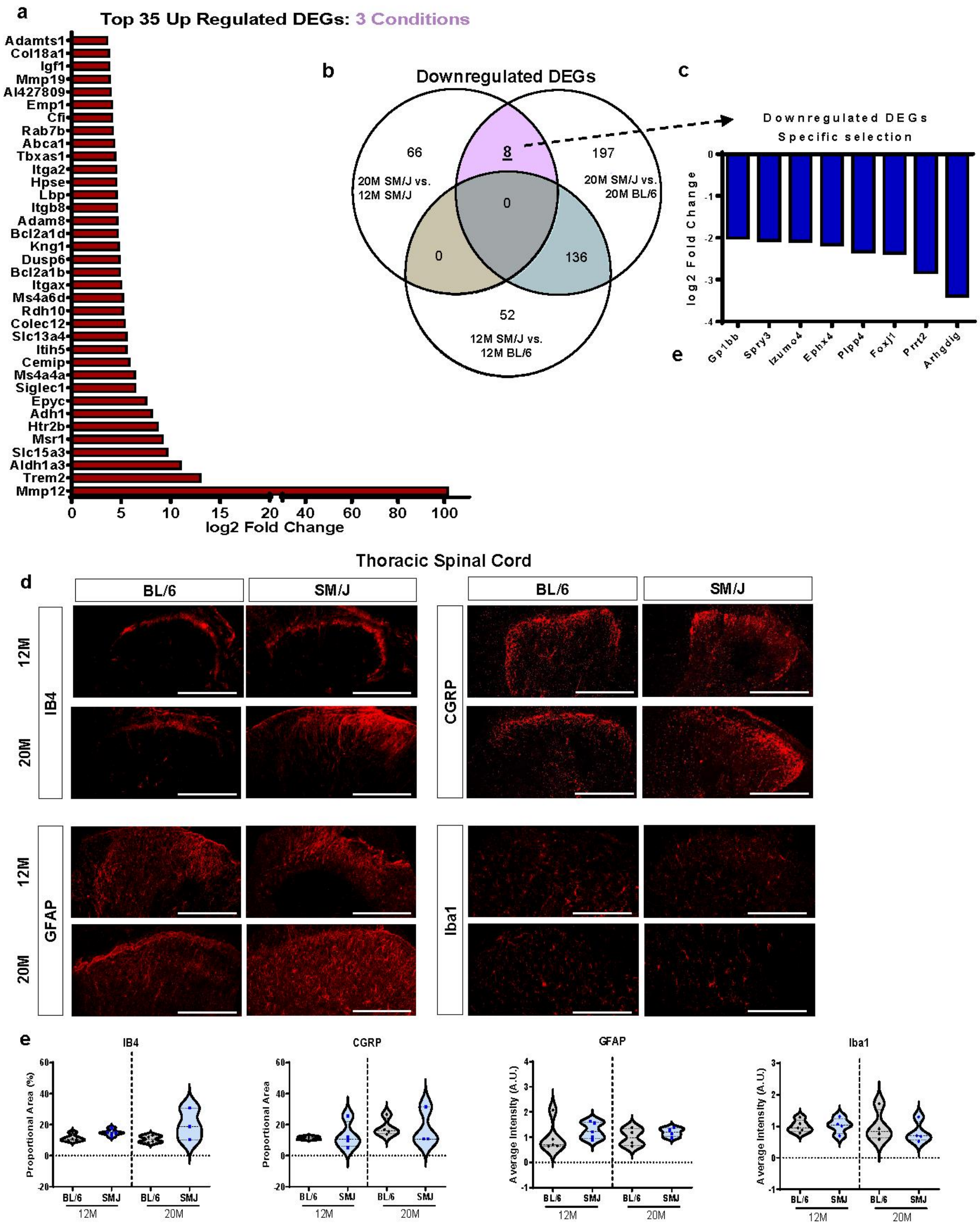

**Supplementary Figure 8:**

**(a)** Top 35 DEGs identified in Figure 4e, based on common upregulated DEGs in 20M SM/J vs. 12M SM/J and 20M SM/J vs. 20M BL/6, but not present in 12M SM/J vs. 12M BL/6 comparison **(b)** Venn Diagram of common downregulated DEGs in 20M SM/J vs. 12M SM/J and 20M SM/J vs. 20M BL/6, but not present in 12M SM/J vs. 12M BL/6 comparison. **(c)** 8 DEGs identified from analysis presented in Suppl. Fig. 8b (highlighted in pink). **(d-e)** Quantitative immunohistochemistry on dorsal horn of thoracic spinal cord, shows no age- or strain-based differences in IB4, CGRP, GFAP, or IBA1 staining and localization. $n_{BL/6\ 12M}$ = 6, $n_{BL/6\ 20M}$ = 4; $n_{SM/J\ 12M}$= 6, $n_{SM/J\ 20M}$ = 4, 2 levels per animal. Two-tailed t-test or Mann-Whitney test was used as appropriate. Data are represented as mean ± SEM. Scale bar A-H' = 200 µm.

Supplementary Figure 9

**a**

| | BL/6 12M | BL/620M | SM/J 12M | SM/J 20M |
|---|---|---|---|---|
| B Cells | 2035 | 1021 | 1361 | 1583 |
| Dendritic Cells | 3 | 7 | 3 | 2 |
| Endothelial Cells | 4 | 5 | 1 | 1 |
| Erythrocytes | 3 | 2 | 14 | 5 |
| Fibroblasts | 0 | 0 | 0 | 2 |
| Granulocytes | 102 | 134 | 27 | 24 |
| Macrophages | 94 | 108 | 65 | 387 |
| Monocytes | 628 | 1208 | 510 | 287 |
| NK Cells | 151 | 96 | 68 | 57 |
| T Cells | 408 | 528 | 361 | 233 |

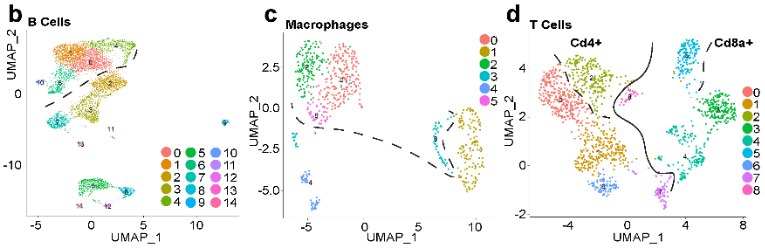


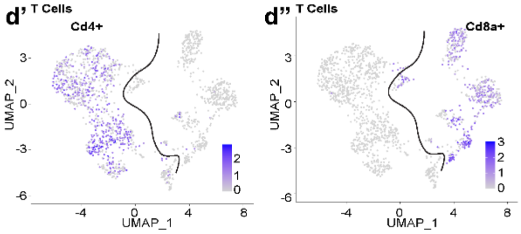


**d'''**

| T Cell Proportion | Cd4+ | Cd8a+ |
|---|---|---|
| BL/6 12M | 20.59% | 26.47% |
| BL/6 20M | 33.33% | 8.33% |
| SM/J 12M | 25.76% | 15.51% |
| SM/J 20M | 31.76% | 10.30% |

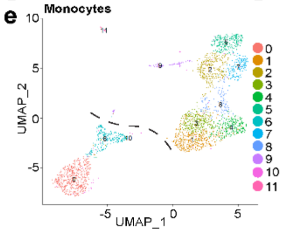


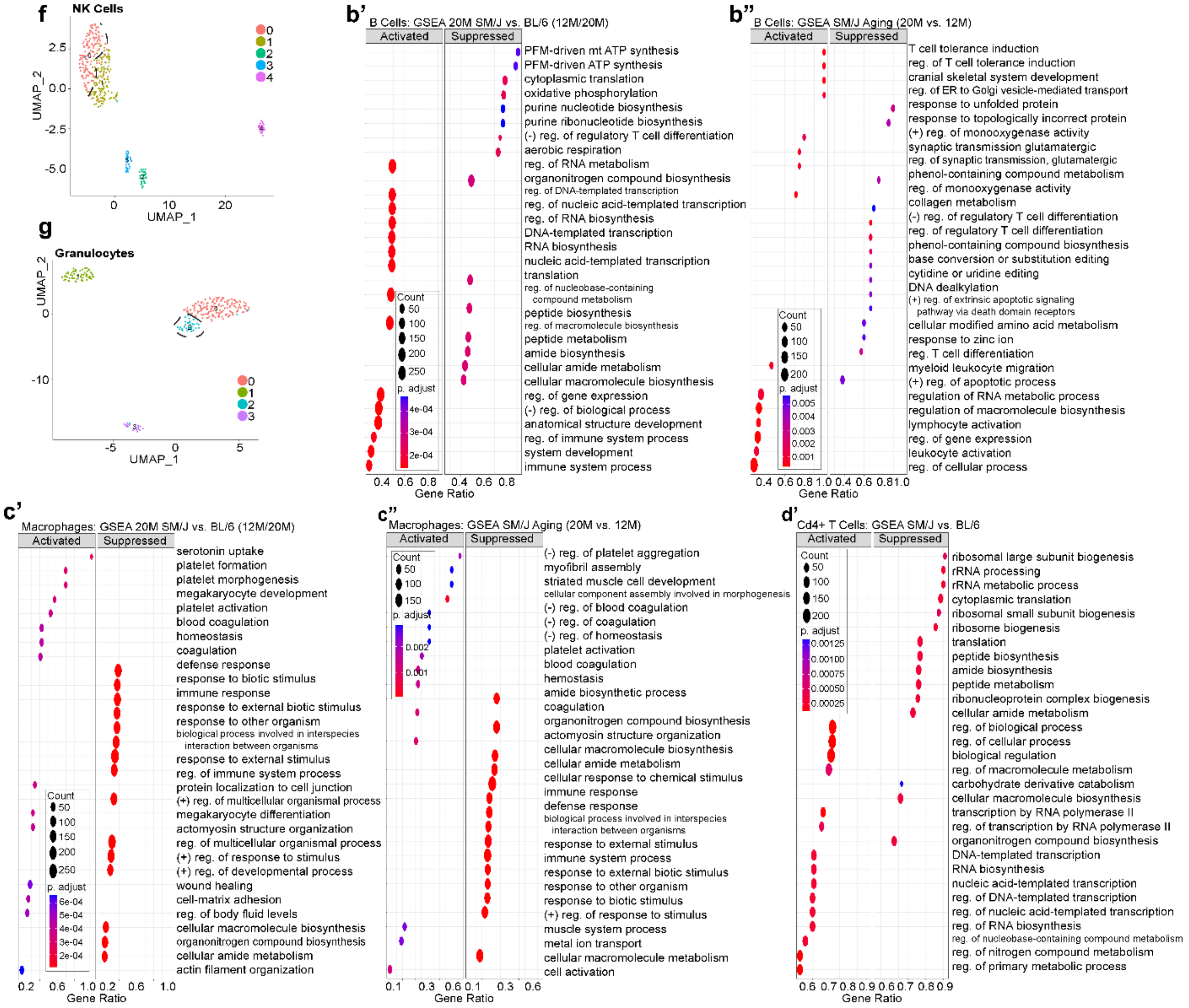

**Supplementary Figure 9:**

**(a)** Cell counts of each cell type identified in scRNA-seq of 12M and 20M BL/6 and SM/J PBMCs **(c-g)** UMAPs labeled according to unbiased subset clustering of **(b)**B Cells, **(c)** Macrophages, **(d)** T cells, (**e)** Monocytes, **(f)** NK Cells, and **(g)** Granulocytes. Gene set enrichment analysis (GSEA) in B cells showing **(b')** 20M SM/J compared to the BL/6 strain at 12M and 20M and **(b")** 20M SM/J compared to 12M SM/J. GSEA in Macrophages showing **(c')** 20M SM/J compared to the BL/6 strain at 12M and 20M and **(c")** 20M SM/J compared to 12M SM/J. **(d')** GSEA showing strain-based differences in CD4+ T cells in SM/J and BL/6 mice. (n=5000 cells/strain/timepoint)

Supplementary Figure 10

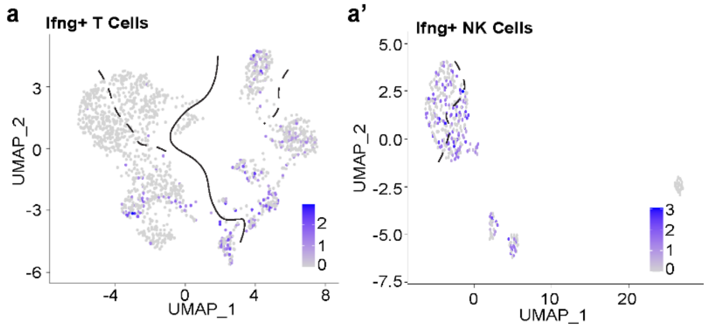


a”

| Ifng+ Proportion | T Cells | NK Cells |
|---|---|---|
| BL/6 12M | 13.73% | 36.42% |
| BL/6 20M | 8.33% | 29.17% |
| SM/J 12M | 4.71% | 25.00% |
| SM/J 20M | 6.87% | 33.33% |

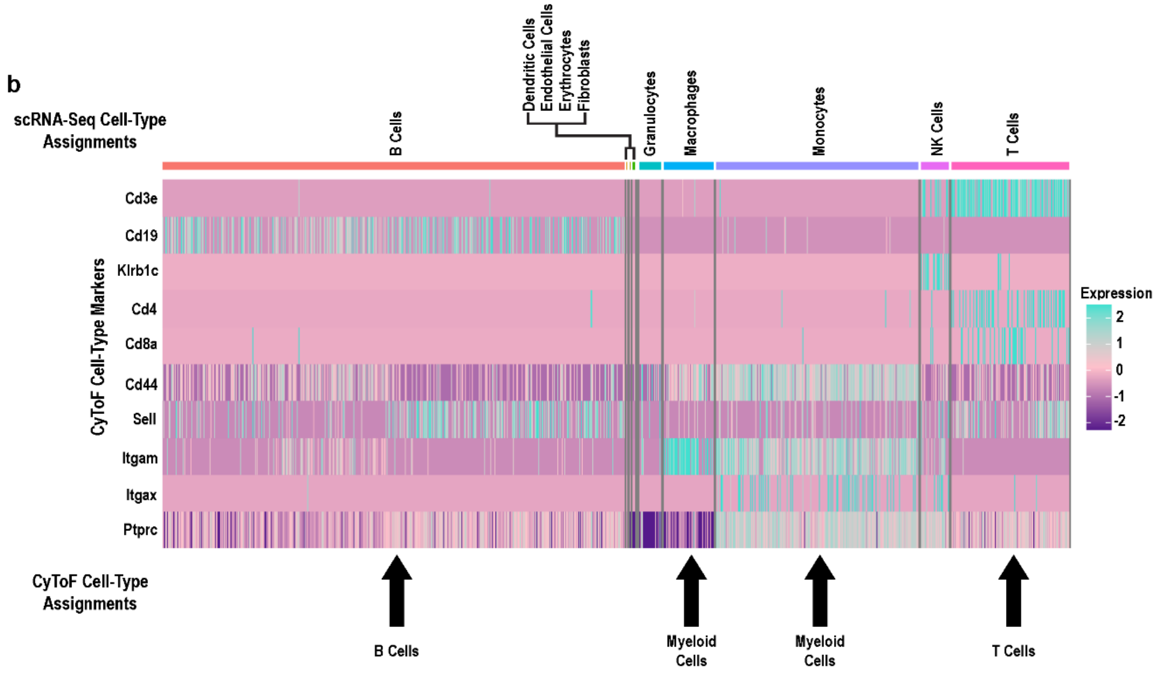


**Supplementary Figure 10: (a)** Ifng+ PBMCs T cells and **(a’)** NK cells, **(a”)** with the proportion of Ifng+ cells, according to strain, age, and cell type. **(b)** PBMC scRNA-seq data labeled according to SingleR cell assignments and organized according to the CyTOF cell labeling criteria.

Supplementary Figure 11

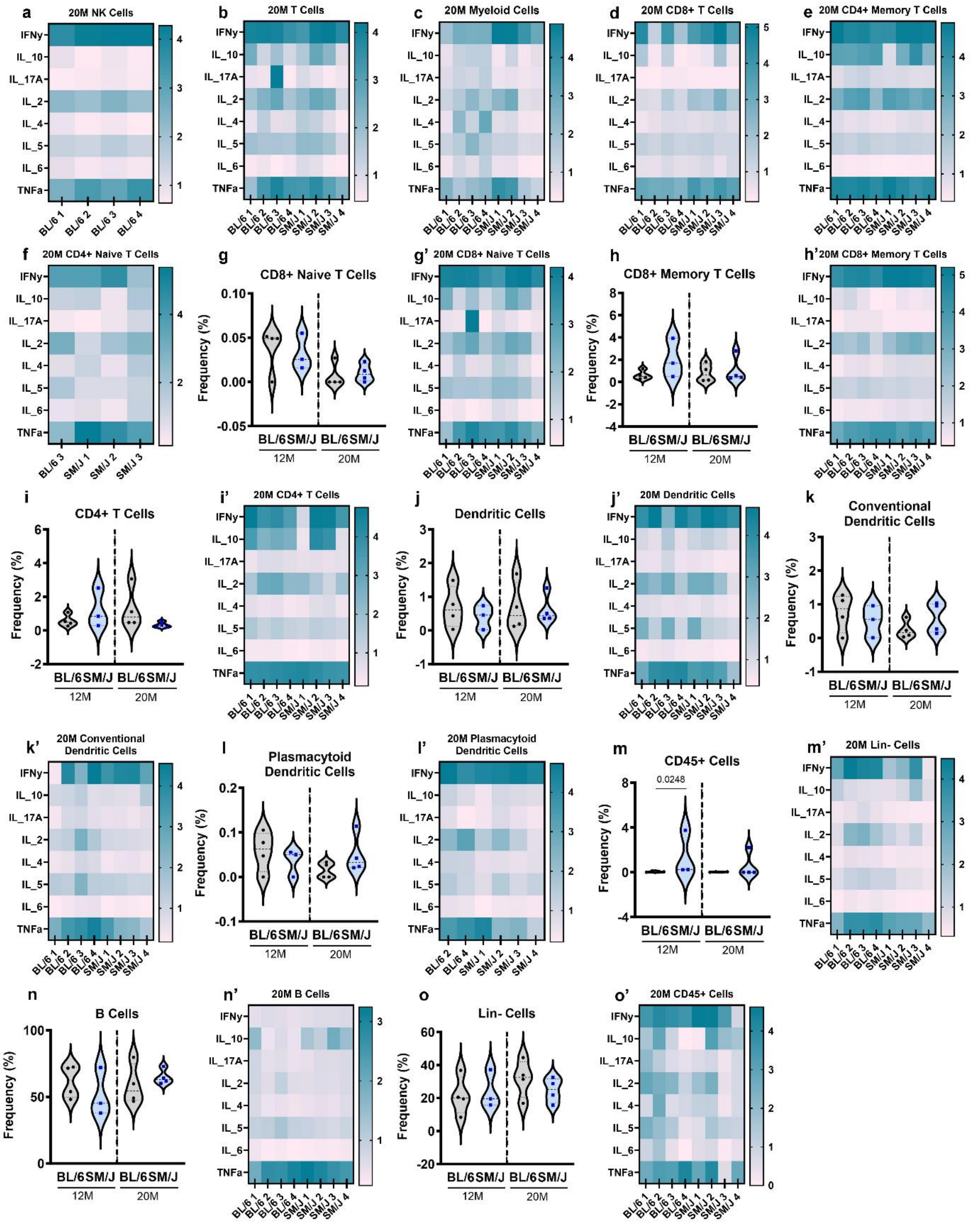

**Supplementary Figure 11:**

**(a-o')** intracellular abundance heatmaps and frequency plots for all splenocyte cell populations which did not demonstrate significance in the CyTOF analysis. ($n_{12MBL/6} = 4$, $n_{12MSM/J} = 3$, $n_{20M\ BL/6} = 4$, $n_{20MSM/J} = 4$) two-tailed t-test or Mann-Whitney test was used as appropriate. Data are represented as mean ± SD.
